# Pathogenetic mechanisms of muscle-specific ribosomes in dilated cardiomyopathy

**DOI:** 10.1101/2025.01.02.630345

**Authors:** Michael R. Murphy, Mythily Ganapathi, Teresa M. Lee, Joshua M. Fisher, Megha V. Patel, Parul Jayakar, Amanda Buchanan, Alyssa L Rippert, Rebecca C. Ahrens-Nicklas, Divya Nair, Rajesh K. Soni, Yue Yin, Feiyue Yang, Muredach P. Reilly, Wendy K. Chung, Xuebing Wu

## Abstract

The heart employs a specialized ribosome in its muscle cells to translate genetic information into proteins, a fundamental adaptation with an elusive physiological role^1–3^. Its significance is underscored by the discovery of neonatal patients suffering from often fatal heart failure caused by severe dilated cardiomyopathy when both copies of the gene *RPL3L* are mutated^4–9^. RPL3L is a muscle-specific paralog^1–3^ of the ubiquitous ribosomal protein L3 (RPL3), which makes the closest contact of any protein to the ribosome’s RNA-based catalytic center^10^. *RPL3L*-linked heart failure represents the only known human disease associated with tissue-specific ribosomes, yet the underlying pathogenetic mechanisms remain poorly understood. Intriguingly, disease is linked to a large number of mostly missense variants in *RPL3L*, and *RPL3L*-knockout resulted in no severe heart defect in either human or mice^3, 11–13^, challenging the prevailing view that autosomal recessive diseases are caused by loss-of-function mutations. Here, we report three new cases of *RPL3L*-linked severe neonatal heart failure and present a unifying pathogenetic mechanism by which a large number of variants in the muscle-specific ribosome led to disease. Specifically, affected families often carry one of two recurrent toxic gain-of-function variants alongside a family-specific putative loss-of-function variant. While the non-recurrent variants often trigger partial compensation of *RPL3* similar to *Rpl3l*-knockout mice, both recurrent variants exhibit increased affinity for the RPL3/RPL3L chaperone GRWD1^14–16^ and 60S biogenesis factors, sequester 28S rRNA in the nucleus, disrupt ribosome biogenesis, and trigger severe cellular toxicity that extends beyond the loss of ribosomes. These findings provide critical insights for genetic screening and therapeutic development of neonatal heart failure. Our results suggest that gain-of-toxicity mechanisms may be more prevalent in autosomal recessive diseases, and a combination of gain-of-toxicity and loss-of-function mechanisms could underlie many diseases involving genes with paralogs.

## Main

The heart is a remarkable organ, beating continuously throughout a human lifetime with a minimal need to replenish its muscle cells (cardiomyocytes)^17^. This extraordinary durability is attributed to many unique adaptations in cardiomyocytes. One of the most fundamental yet least understood adaptations is that cardiomyocytes use a specialized ribosome to decode genetic information and synthesize proteins^1–3^. Although ribosomes in all cells use the same genetic code, the composition of the ribosome itself—a delicate molecular machine comprising 80 ribosomal proteins and 4 ribosomal RNAs (rRNAs)—can differ across cell types^18–24^. The most striking variation occurs in cardiomyocytes. During vertebrate heart development, the ubiquitous core ribosomal protein L3 (RPL3)—a gatekeeper of the ribosome’s decoding center^25^ and the protein closest to the ribosome’s RNA-based active site^10^— is gradually replaced by its paralog, RPL3-like (RPL3L)^1, 26^. RPL3L shares 74% amino acid identity with RPL3^1, 2^ and eventually becomes the dominant form in adult human left ventricular cardiomyocytes (95%)^27^, to a lesser extent in skeletal muscle (60%), and barely detectable in non-muscle cells^28^. Interestingly, this switch is reversed during adult muscle growth (hypertrophy) following injury or atrophy^2, 12, 29^. The physiological role of this highly conserved and tightly regulated ribosomal switching remains poorly understood, highlighting a crucial gap in our understanding of cardiac biology at the most fundamental level.

The essential role of RPL3L and the specialized ribosome in the heart is underscored by an increasing number of severe neonatal heart failure cases caused by dilated cardiomyopathy (DCM) associated with *RPL3L* mutations^4–9^. First identified by our team^4^, this condition has been classified as a new subtype of DCM, Dilated Cardiomyopathy-2D (CMD2D, OMIM# 619371). It stands out as most known DCM genes encode proteins associated with the sarcomere^30–32^, the contractile unit of the muscle. *RPL3L*-linked DCM was the first—and remains the only—known human disease caused by mutations in tissue-specific ribosomes. Among seven independent affected families, individuals with mutations in both *RPL3L* alleles consistently develop severe DCM, often resulting in fatal heart failure^4–9^. In contrast, heterozygous parents and siblings carrying a single mutant allele remain unaffected with the early onset DCM. While the autosomal recessive inheritance pattern suggests a loss-of-function of these variants and RPL3L deficiency as the likely cause of DCM, *Rpl3l* knockout (KO) mice generated by multiple labs display only mild cardiac dysfunction^3, 11–13^. This is unexpected given the 96% homology between human and mouse RPL3L proteins and that the highly regulated RPL3-RPL3L switch is also conserved in mice^12^.

The discrepancy between early onset fatal disease in humans and mild phenotypes in mice suggests either species-specific differences in heart biology or that the DCM-causing autosomal recessive *RPL3L* mutations have effects beyond simple loss-of-function, rendering them unsuitable for modeling with KO mice. While rarely reported, autosomal recessive mutations can cause disease via toxic gain-of-function^33^. Notably, in all *Rpl3l* KO mouse models, regardless how they were generated—whether through a frameshifting deletion of the entire exon 2^3^, short frameshifting deletions in exon 5^11, 12^, or poly(A) insertion in exon 1^13^—there is a consistent up-regulation of the ubiquitous paralog Rpl3. This compensatory increase in Rpl3 has been proposed to explain the absence of a severe cardiac phenotype in mice, as Rpl3 is expected to largely fulfill the functional role of Rpl3l. The molecular mechanism underlying the *Rpl3* compensation remains unclear. Whether a lack of *RPL3* compensation in human underlie the severe DCM symptoms remains unexplored. A deeper understanding of the pathogenetic mechanism of *RPL3L*-linked DCM, including whether *RPL3* compensation occurs and how to activate the compensation, may inform future management and the development of new therapeutics for these patients. This is significant as nearly half of all DCM cases—and two-thirds of pediatric cases—remain of unknown etiology^30, 31, 34, 35^. More broadly, this lethal disease provides a rare opportunity to explore the physiological roles of ribosomes, their tissue-specific functional specializations, and the mechanisms by which their dysfunction contributes to human disease.

In this study, we present three newly identified cases of *RPL3L*-linked severe neonatal heart failure and uncover a distinctive pathogenetic mechanism through integrated analyses of population genetic data, patient cardiac tissue, and isogenic cells expressing *RPL3L* variants.

### Hotspot G27D and D308N/V are potential driver mutations

We identified three additional unrelated cases of *RPL3L*-associated early onset CMD2D, increasing the total number of affected families to ten (Supplemental Information). Similar to previously reported cases^4–9^, all three families are associated with compound heterozygous variants, i.e., each affected individual carry two different variants on separate alleles. Interestingly, all three cases share the D308N variant, a missense amino acid substitution at position 308 that changes aspartic acid (D) to asparagine (N). The affected individuals co-inherit one of three additional variants: a missense variant (R58Q), a truncating frameshift variant (T340Nfs*25), or a splice donor variant (c.1167+1G>A) that causes skipping of exon 9 (Δexon9)^36^. The latter two variants are predicted to result in loss-of-function.

The three new cases highlight three prominent features characteristic of *RPL3L*-linked DCM across all reported cases: compound heterozygosity, hotspot variants, and the pairing of hotspot variants with putative loss-of-function variants. First, all but one affected families exhibit compound heterozygous variants in *RPL3L* (lines in **Fig. 1a**). The only exception is a homozygous R116H variant identified in a consanguineous population^7^. Across ten families, at least one unique variant is present per family, with a total of 14 distinct variants identified. Second, while variants are distributed across nearly the entire RPL3L protein (**Fig. 1a-b**), two notable hotspots emerge: D308 (D308N or D308V), observed in five families, and G27 (G27D), found in three additional families (**Fig. 1a**). Collectively, D308N/V and G27D account for 80% (8/10) of *RPL3L*-linked DCM families. These families appear to be unrelated, as evidenced by their diverse geographic locations, with the three families carrying the G27D variant residing in the United States, Spain, and China. Lastly, among the eight families carrying one of the two hotspot variants, four harbor a paired frameshift or splice variant (**Fig. 1a**), typically considered (partial) loss-of-function, raising the possibility that the missense variants identified in the remaining four families may also result in loss of function.

**Figure 1.**
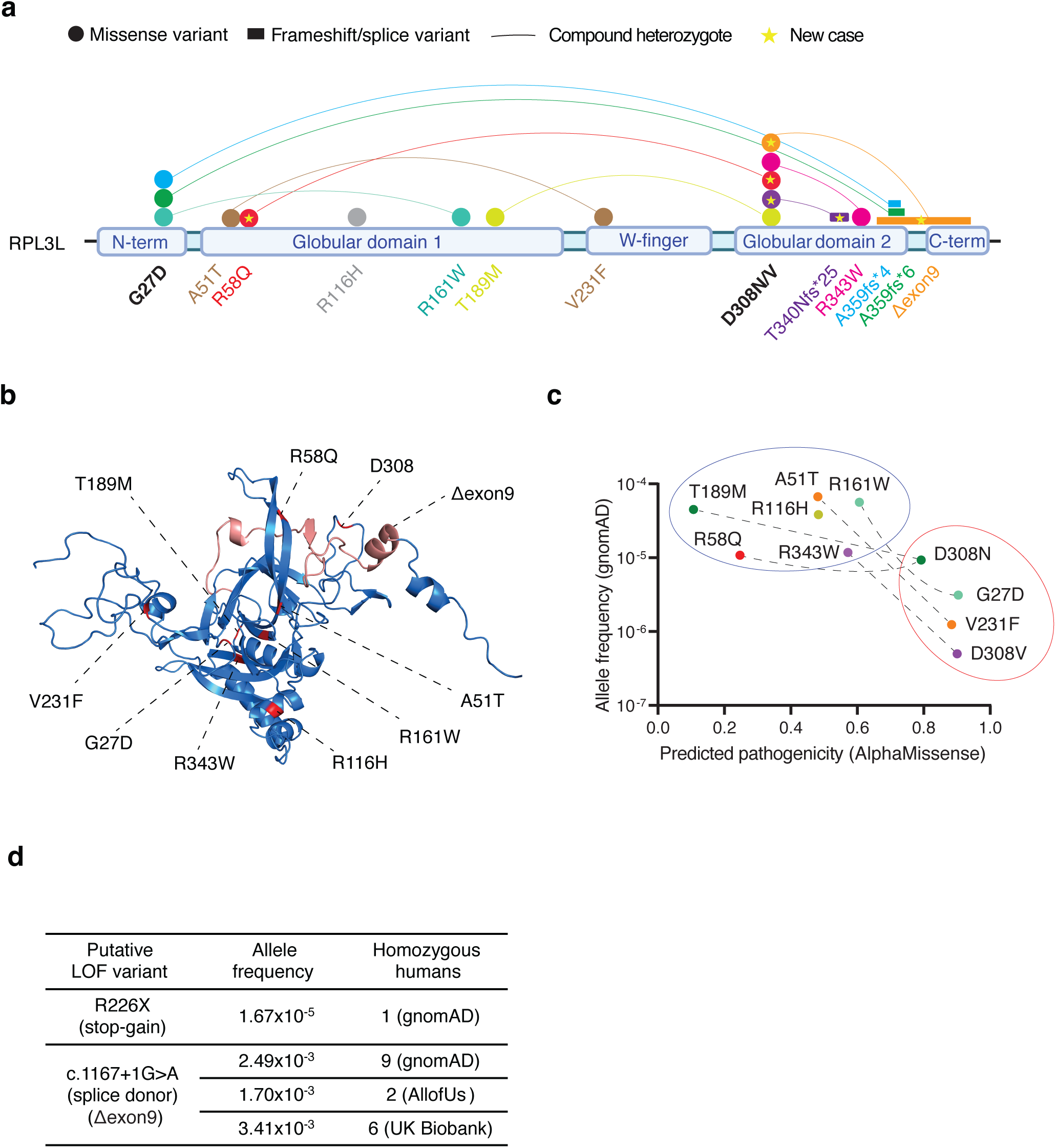
Hotspot variants G27D and D308N/V as potential drivers in *RPL3L*-linked DCM. **a**, Schematic representation of neonatal DCM-associated variants across the RPL3L protein. Each circle denotes a missense variant, while rectangles represent frameshift variants or the splice variant (Δexon9). Variants are color-coded by family, with lines connecting compound heterozygous variants identified in the same family. Variants identified in the three new cases are highlighted with a star. The protein domains within RPL3L are labeled. The hotspot variants G27D and D308N/V are emphasized in bold. **b**, Mapping of the variants (in red) onto the AlphaFold-predicted 3D structure of RPL3L. The region encoded by exon 9, deleted in the splice variant (Δexon9) is highlighted in salmon. Frameshift variants are not shown. **c**, Clustering of RPL3L missense variants based on allele frequency (gnomAD) and predicted pathogenicity (AlphaMissense). Variants identified in the same family are connected by dashed lines. **d**, Putative human *RPL3L* KO in the general population. LOF: loss-of-function.

Notably, the ten missense variants can be categorized into two groups based on allele frequency and pathogenicity predicted by AlphaMissense^37^, a state-of-the-art deep learning model built on the AlphaFold2 protein structure prediction tool^38^ (**Fig. 1c**). The hotspot variants G27D and D308N/V fall into the group with extremely low allele frequencies in the general population (<10^-5^, gnomAD^39^) and are predicted to be highly pathogenic. The rarity of these hotspot variants in the general population argues against the hypothesis that their recurrence in patients is due to increased DNA mutability. Instead, the repeated occurrence of these ultra-rare missense mutations in a significant proportion of DCM families strongly challenges a loss-of-function mechanism for these variants, especially considering that neither G27 nor D308 corresponds to residues known to be critical for RPL3/RPL3L’s role in the ribosome^10, 25^. The high-pathogenicity group also includes the V231F variant, another extremely rare (∼10^-6^) variant in the population and predicted to be highly pathogenic by AlphaMissense. It is possible that with more cases to be discovered, V231F may also be found to be a hotspot mutation similar to G27D and D308N/V in the same group. Variants in the low-pathogenicity group are more frequent (still very rare, 10^-5^ to 10^-4^) and predicted to be less pathogenic (**Fig. 1c**). Interestingly, with the exception of the homozygous R116H mutation, each affected individual carries one mutation from the high-pathogenicity group and another mutation from the low-pathogenicity group (linked by dashed lines).

Consistent with the absence of a severe cardiac phenotype in *Rpl3l* KO mice, live human knockouts of *RPL3L* have been identified in large-scale genetic studies, including gnomAD^39^, All of Us^40^, and UK Biobank^41^ (**Fig. 1d**). One individual carries a premature stop codon at arginine 226 in both alleles (R226X), truncating nearly half of the protein (full-length RPL3L is 407 amino acids), a mutation almost certainly resulting in loss-of-function. Additionally, a total of 17 individuals are homozygous for the same splice donor variant found in one of the DCM families (c.1167+1G>A). As shown previously^36^, the splice variant causes exon 9 deletion, removing a highly conserved 40- amino-acid region in RPL3L (salmon in **Fig. 1b**). Although this splice donor variant is associated with an increased risk of atrial fibrillation in multiple association studies^36^, homozygous individuals carrying this variant appear to be alive and do not exhibit severe early-onset DCM. There are also nine putative knockout humans in the Regeneron Genetics Center Million Exome (RGC-ME) database^42^ although detailed information about these variants is not available. The lack of severe cardiac dysfunction in both putative *RPL3L* KO humans and *Rpl3l* KO mice strongly suggests that the missense *RPL3L* mutations identified in DCM patients are not simply loss-of-function variants.

In summary, our meta-analysis of *RPL3L*-linked DCM cases reveals that affected individuals frequently carry a recurrent, highly pathogenic missense variant paired with either a frameshift loss-of-function variant or a low-pathogenicity missense variant. Combined with the absence of severe heart defects in *RPL3L*-knockout mice and humans, these findings suggest that *RPL3L*-linked DCM is likely driven by a mechanism other than loss-of-function, despite its autosomal recessive inheritance pattern.

### Ribosome defects in explanted patient hearts

To investigate the pathogenic mechanisms of *RPL3L* mutations in DCM, we initially focused on a patient carrying the hotspot D308N variant and the T189M missense variant. This patient, diagnosed and treated at our center, underwent heart transplantation and is alive to date^4^. Using explanted heart ventricular tissues, we extracted total RNA to perform RNA-seq and compared the results to age-matched control healthy heart ventricular tissues.

Unexpectedly, we observed an abnormality when the total RNA was subjected to Bioanalyzer electrophoresis for quality check (**Fig. 2a**). Specifically, there is a drastic reduction in the 28S rRNA peak relative to the 18S peak. The 28S rRNA is the scaffold of the 60S large subunit of the ribosome. RPL3L (and the paralog RPL3) is a core component of the 60S subunit and plays a vital role in both 28S rRNA biogenesis and pre-60S ribosome assembly via direct binding to the 28S rRNA^43^. The loss of 28S rRNA indicates a defect in 28S/60S biogenesis or turnover in patient cells, presumably due to the D308N/T189M mutations in *RPL3L*.

**Figure 2.**
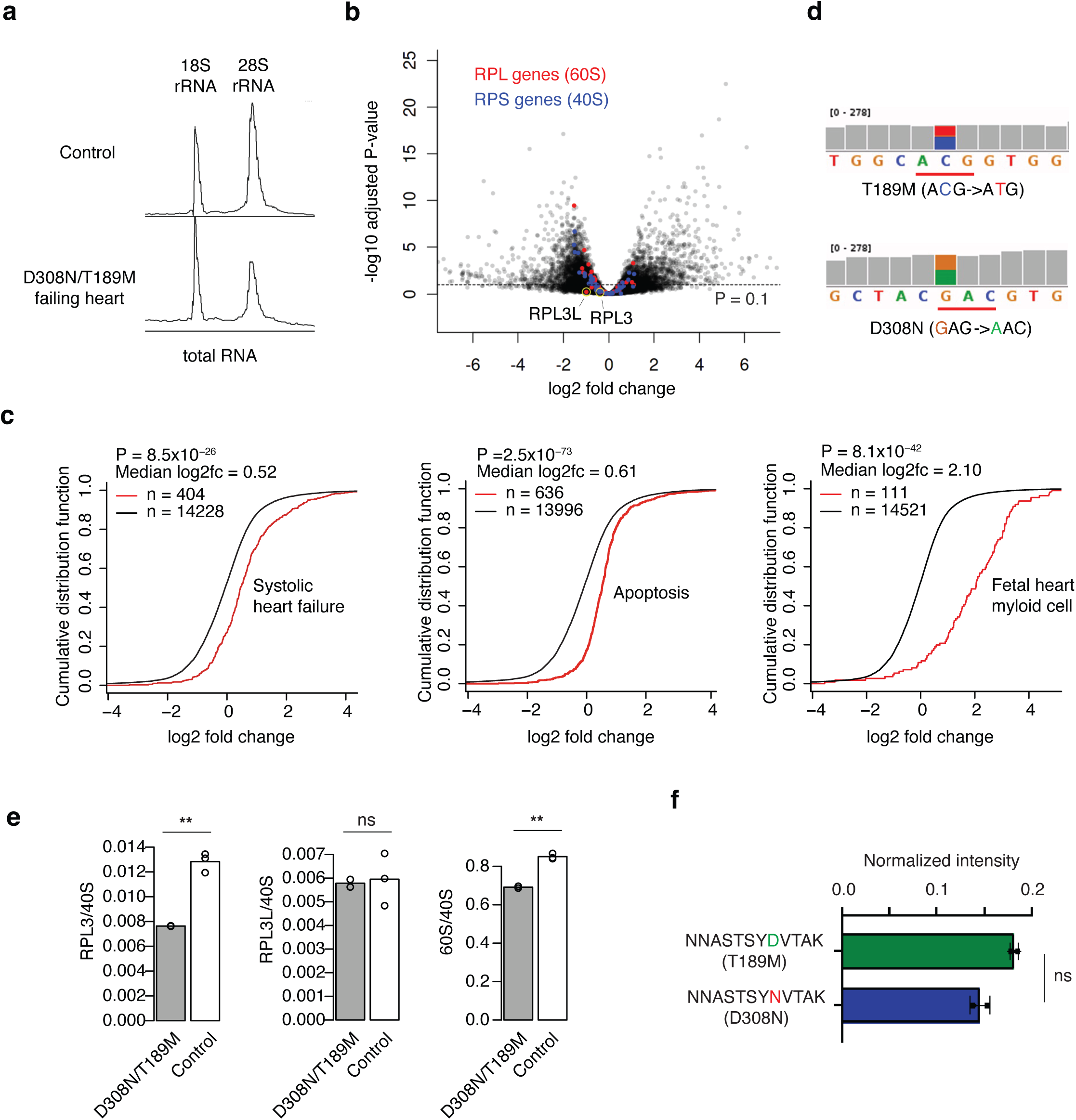
Ribosome defects in explanted patient hearts. **a**, Bioanalyzer electrophoresis of total RNA isolated from explanted patient heart ventricular tissue and a control sample (human AC16 cells). **b**, Differential gene expression analysis comparing explanted heart tissue (N=2)and age-matched healthy control heart ventricular tissue (N=3). Genes encoding 60S or 40S ribosomal proteins are colored red or blue, respectively. *RPL3* and *RPL3L* are highlighted by yellow circles. **c**, Cumulative distribution function plots for three significant gene signatures. Red indicates genes of interest and black indicates all other genes. The median log2 fold change (log2fc) and Kolmogorov–Smirnov test P values are also shown. **d**, RNA-seq read coverage (BAM file visualized in IGV genome browser) showing that both D308N and T189M alleles are equally expressed. **e**, Mass spectrometry quantification of RPL3 (left), RPL3L (center), and total 60S proteins (right) relative total 40S proteins in D308N/T189M patient ventricular tissue (N=2) and age-matched control tissue (N=3). ** P < 0.0001. **f**, Intensity of the peptide containing the D308 residue in the T189M variant and the peptide containing the N308 residue in the D308N variant normalized to the total intensity in RPL3L. ns: not significant.

The RNA-seq data revealed widespread alterations in gene expression, including 1,639 significantly up-regulated genes and 2,361 significantly down-regulated genes (**Fig. 2b**). Pathway analyses confirmed up-regulation of gene signatures associated with systolic heart failure and apoptosis, and interestingly, a massive up-regulation of genes associated with fetal cardiac myeloid cells, consistent with increased inflammation in failing hearts^44^ (**Fig. 2c**).

In line with impaired ribosome biogenesis, many ribosomal protein genes are differentially expressed, with more down-regulated than up-regulated (**Fig. 2b**). *RPL3L* is down-regulated by approximately two-fold, although not significantly (adjusted P = 0.6). Both D308N and T189M alleles are expressed at a similar level as indicated by sequencing read coverage (**Fig. 2d**).

Surprisingly, unlike the robust increase of *Rpl3* mRNA and protein in *Rpl3l* KO mice, there was no compensatory up-regulation of *RPL3* mRNA in the patient’s heart tissue (**Fig. 2b**). Instead, *RPL3* is slightly down-regulated (by 24%, adjusted P=0.66) like many other ribosomal protein genes.

To confirm the lack of *RPL3* compensation in the DCM heart tissue, we also examined protein expression using mass spectrometry. Indeed, when normalized to the total abundance of 40S ribosomal proteins, there is a significant decrease in RPL3 abundance with no change in RPL3L (**Fig. 2e**). Consistent with a 60S biogenesis defect, there is a mild but significant decrease in total 60S ribosomal protein level relative to that of the 40S. Similar to the RNA-seq data (**Fig. 2d**), the D308N variant is expressed at a comparable level as the T189M variant (**Fig. 2f**).

In the absence of compensatory RPL3 up-regulation, there will be a shortage of functional 60S ribosomes in patient cells, as neither functional RPL3L nor RPL3 is present. Given the critical role of ribosomes in cellular function, the loss of functional ribosomes likely contributes to cardiomyocyte death and the progression of DCM. While caveats remain (see Discussion), the lack of RPL3 compensation observed in human DCM patients underscores a critical divergence from the *Rpl3l* KO mouse model and suggests that at least one of the two missense *RPL3L* variants (D308N and T189M) are unlikely to function as simple loss-of-function variants.

In summary, analysis of explanted heart tissues from DCM patients demonstrates a defect in 60S ribosome biogenesis and a lack of RPL3 compensation. These defects likely contribute to the loss of functional ribosomes, cardiomyocyte death, and the development of heart failure in *RPL3L*-linked DCM.

### Hotspot variants gain toxic function to drive ribosome biogenesis defects and cellular toxicity

It remains unclear whether the ribosome biogenesis defect observed in explanted patient tissue is caused by the hotspot D308N mutation or the accompanying T189M variant. Both variants are expressed at comparable levels with no significant allele imbalance (**Fig. 2d** and **Fig. 2F**).

To determine the individual impact of these variants, we generated isogenic cell lines expressing various RPL3L variants in the absence of RPL3 (**Fig. 3a**). These cell lines were derived from human AC16 cells, ventricular cardiomyocyte-like cells commonly used to study cardiac gene expression and function^45^, including the regulation of ribosomes and translation in cardiac hypertrophy^46^. While AC16 cells do not fully recapitulate primary cardiomyocytes, our results described below demonstrate their suitability for studying ribosome-related processes. Of note, *RPL3L* is not expressed in iPSC-derived cardiomyocytes (iPSC-CMs) (**Extended Data Fig. 1**), consistent with the notion that iPSC-CMs resemble immature fetal cardiomyocytes^47, 48^, while *RPL3L* expression is restricted to mature cardiomyocytes.

**Figure 3.**
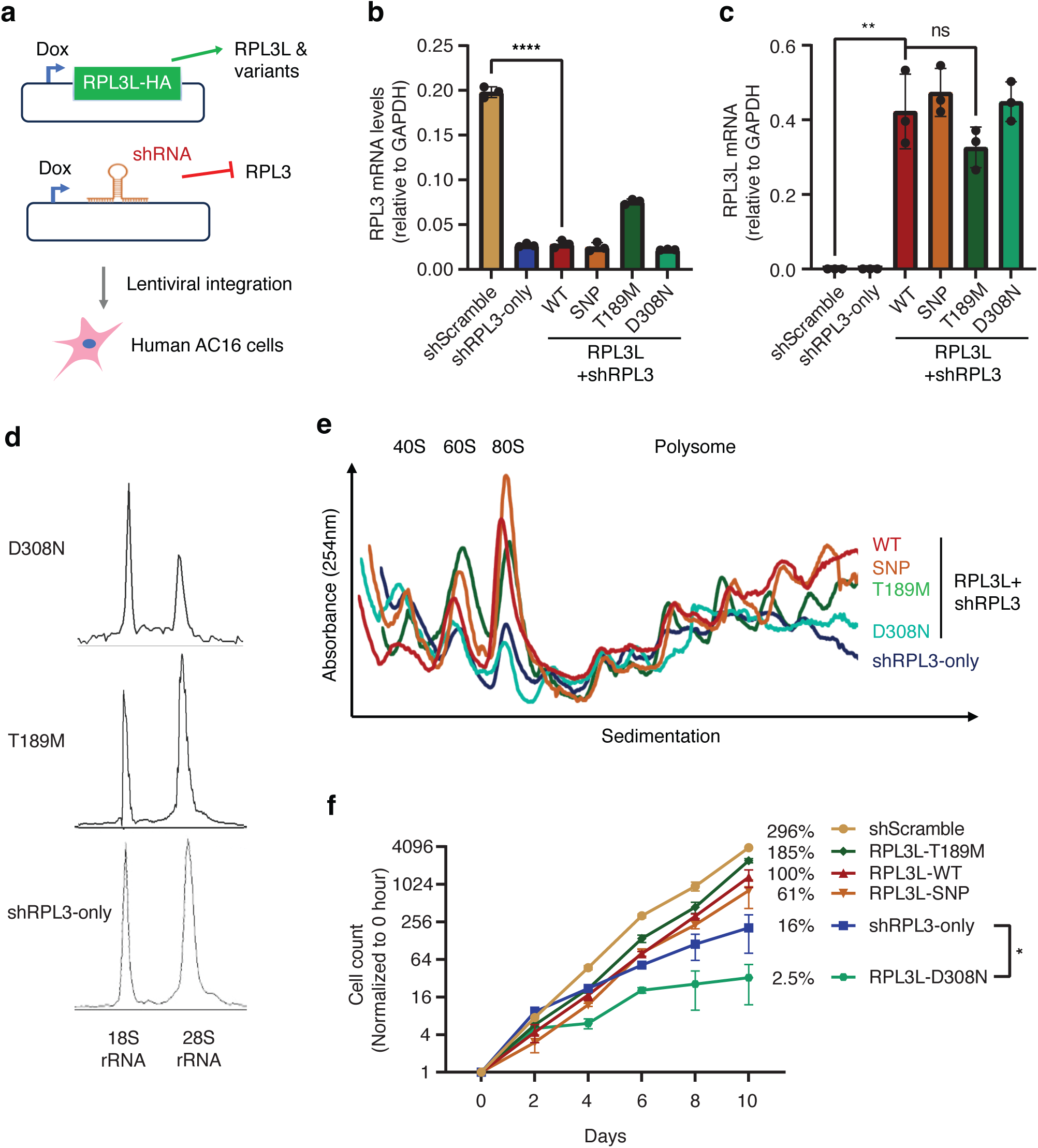
Hotspot variants disrupt ribosome biogenesis and impairs cell viability. **a**, Isogenic AC16 human cardiomyocyte-like cell lines were generated to simultaneously knockdown (KD) RPL3 and overexpress HA-tagged RPL3L (or its variants) via a dox-inducible promoter. **b**, RPL3 knockdown after dox treatment. Total RNA was isolated from indicated cell lines and analyzed for RPL3 levels. shScramble: cells expressing only a control scrambled shRNA. shRPL3: cells expressing only the RPL3 targeting shRNA without RPL3L or its variants. RPL3L-WT/SNP/T189M/D308N: cells with RPL3 KD and expresses either wild type or the indicated variant of RPL3L. **c**, Efficient expression of RPL3L variants in AC16 cells after 120 hours Dox treatment. Total RNA isolated from indicated cells was analyzed for RPL3L levels (N = 3 independent experiments). **d**, Bioanalyzer electrophoresis on total RNA from indicated cell lines was quantified by ImageJ. **e**, Loss of translation capacity in shRPL3 and D308N cells, but not control or T189M cells. RPL3+ (shScramble) and RPL3L-expressing cells were separated by ultracentrifugation and fractionated in the presence of cycloheximide (N = 3). **f**, Cell growth defects in D308N and shRPL3 cells. AC16-derived cells were seeded at equal density and counted every 2 days (N = 4). The y-axis is shown on a logarithmic scale. The percentage values represent the cell density at day 10 relative to the RPL3L-WT cell line. **b ,c, f ,** Data are mean ± s.d. *: P < 0.05.

Wild-type AC16 cells naturally express only *RPL3*, not *RPL3L*. To switch to *RPL3L* variants, we first introduced a doxycycline (dox)-inducible shRNA targeting *RPL3* to deplete RPL3 in an inducible manner. This cell line (shRPL3-only) serves as a critical baseline control for the cellular defect and toxicity caused by the loss-of-function of ribosomal proteins. Using shRPL3-only cells, we further integrated a second dox-inducible transgene expressing HA-tagged *RPL3L*, either the wild type or variants (**Fig. 3a**). In addition to DCM-linked *RPL3L* variants, we also included a common SNP (rs34265469) predicted to result in a benign missense variant (P291L). This SNP is the most common missense variant in *RPL3L*, with an allele frequency of 4.26% (TOPMed) and is homozygous in at least 715 living individuals. This isogenic system enabled allele-specific switching from *RPL3* to *RPL3L* at physiological level upon dox treatment (**Fig. 3b-c**).

Using these isogenic cell lines, we found that the loss of the 28S rRNA observed in the explanted patient heart tissue (**Fig. 2a**) was faithfully recapitulated in cells expressing the D308N variant, but not in those expressing the T189M variant (**Fig. 3d**). Notably, the loss of 28S rRNA in D308N cells is much stronger than that of the shRPL3-only cells (**Fig. 3d**). Given that the only difference between these two cell lines is the expression of the D308N variant, the stronger defect indicates toxic gain-of-function of the D308N variant. Additionally, the weaker effect in the shRPL3-only cells suggests that ribosome loss alone is insufficient to fully account for the defects observed in patients (**Fig. 2a**).

Consistent with 60S ribosomal subunit biogenesis defects, sucrose gradient sedimentation revealed a dramatic reduction in 60S but not 40S ribosomal subunits in D308N-expressing cells compared to WT, SNP, or T189M cells (**Fig. 3e**). Accordingly, monosomes (80S) and translating polysomes (poly-ribosomes), both of which contain 60S, are also depleted in D308N-expressing cells (**Fig. 3e**). Notably, the loss of polysomes in D308N-expressing cells is comparable to cells not expressing any *RPL3L* variant (shRPL3-only) (**Fig. 3e**), suggesting the D308N variant of RPL3L almost completely blocks ribosome biogenesis.

In line with a severe loss of ribosomes, D308N-expressing cells exhibited a severe growth defect (**Fig. 3f**, note that the y-axis is on a logarithmic scale). Expressing D308N for 10 days led to a 40- fold reduction in cell counts (compared to RPL3L-WT cells, **Fig. 3f**, green vs. red). Interestingly, cells expressing the D308N variant exhibit a 6.4-fold greater growth defect compared to cells lacking RPL3L expression (shRPL3-only) (green vs. blue, **Fig. 3f**), again suggesting toxic gain-of-function. Of note, the control cell line expressing RPL3 (shScramble) grew faster than cells expressing wild type RPL3L (RPL3L-WT), mirroring a previous mouse study showing that Rpl3l expression is associated with reduced protein synthesis and slower muscle cell growth^2^. Interestingly, cells expressing the T189M variant of *RPL3L* grew faster than cells expressing the wild type *RPL3L*, which may be attributed to a higher expression of *RPL3* in this cell line (**Fig. 3b**, also see below the discussion on *RPL3* compensation). The lack of deleterious effects associated with the T189M variant aligns with its low pathogenicity prediction by AlphaMissense (**Fig. 1c**) and is further supported by the presence of a living homozygous T189M individual in the population genotyped in TOPMed. Taken together, these results indicate that the D308N variant is a toxic gain-of-function variant and the primary driver of the disease in the patient carrying D308N and T189M.

Next, we investigated the other hotspot variant G27D, together with the R161W variant found in the same affected individual^4^. Similar to the D308N variant, cells expressing the G27D variant grew substantially slower than those expressing the wild-type *RPL3L* (33-fold fewer cells by day 10, **Extended Data Fig. 2a**), whereas cells expressing the R161W variant grew slightly faster than cells expressing the WT RPL3L, similar to T189M. Consistent with a growth defect, G27D but not R161W-expressing cells are depleted of 60S and 80S ribosomes as well as polysomes (**Extended Data Fig. 2b**). These findings confirm that the hotspot variants G27D and D308N, but not their co-inherited non-hotspot variants T189M and R161W, are responsible for ribosome biogenesis defects and cellular toxicity.

### Hotspot variants mislocalize and alter interactions with ribosome biogenesis factors

To further explore the molecular basis of the toxic gain-of-function of the hotspot variants, we examined the subcellular localization of RPL3L variants using fluorescence imaging of the HA tag. In contrast to the predominantly cytoplasmic localization of the wild-type RPL3L and the common SNP variant expected from a ribosomal protein, the hotspot variants D308N and G27D both exhibited nearly exclusive nuclear localization (**Fig. 4a**, green, quantified in **Fig. 4b**). Given the role of RPL3/RPL3L in 60S ribosome biogenesis, this mislocalization of the hotspot variants likely disrupt 60S ribosome assembly. Supporting this, fluorescent probes for 28S rRNA revealed a significant decrease in cytoplasmic 28S rRNA levels in D308N or G27D-expressing cells relative to the nucleus (**Fig. 4a**, red, quantified in **Fig. 4c**). Notably, this 28S rRNA localization defect was not observed in cells not expressing any RPL3L variant (shRPL3-only), consistent with a toxic gain-of-function of both hotspot variants.

**Figure 4.**
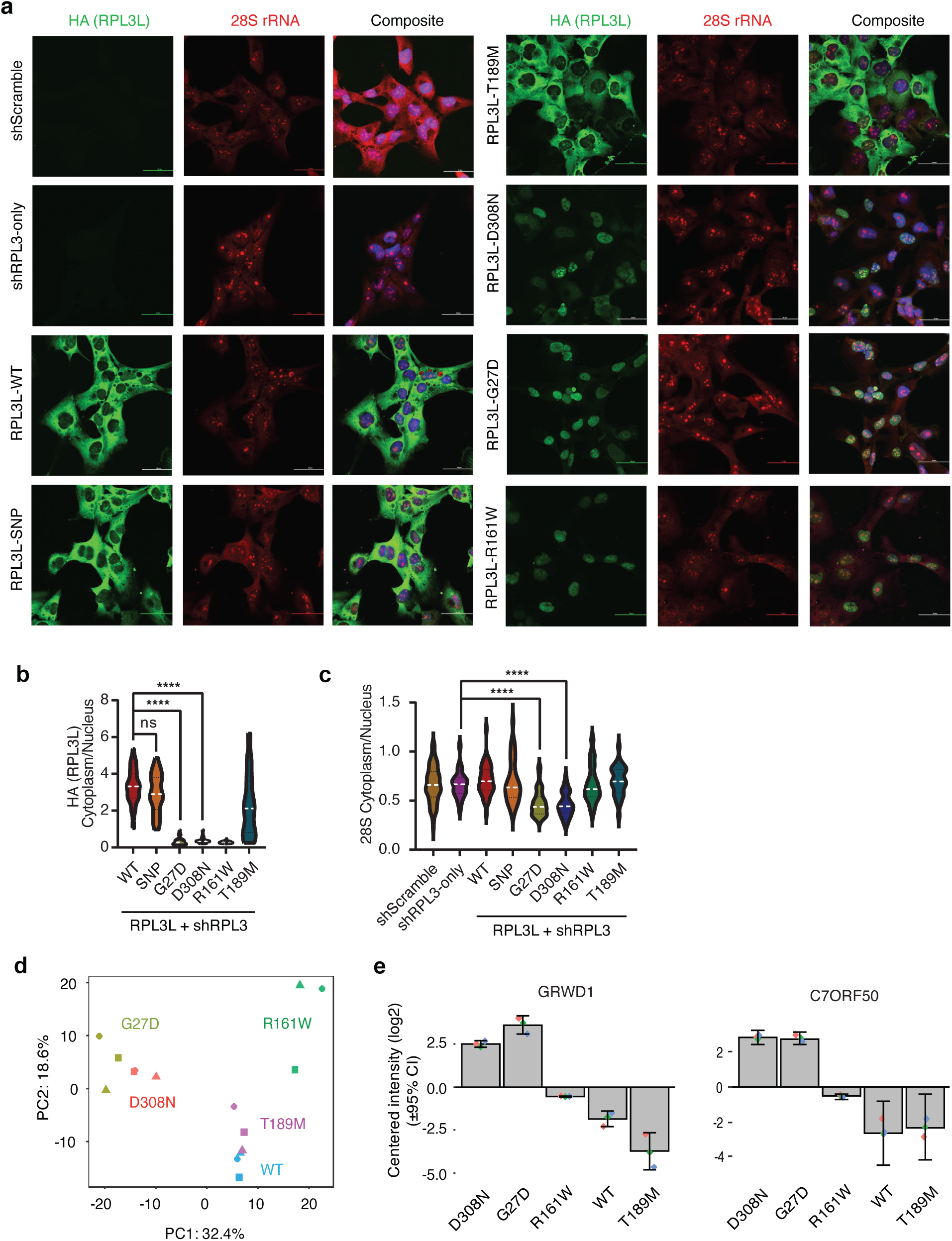
Hotspot variants mislocalize and alter interactions with ribosome biogenesis factors. **a**, Cells were stained for anti-HA antibody (RPL3L and variants) and 28S rRNA FISH probes and analyzed by 60x confocal microscopy. Scale bar = 50μm. **b**, Violin plot quantification of cytoplasmic and nuclear abundances of HA (RPL3L) signals for indicated cell lines, measured per cell (N = 26-41 cells). **c**, same as b but for 28S rRNA. One-way ANOVA was used for statistical analysis. **d**, PCA analysis of the proteins co-immunoprecipitated with RPL3L variants. N=3. **e**, Enrichment of GRWD1 and C7ORF50 in proteins co-immunoprecipitated with the hotspot variants D308N and G27D compared to other variants. Peptide intensities are normalized by the median of all samples and then log2-transformed. Error bars represent standard deviation. ns, not significant; ** P < 0.001; *** P < 0.0001; **** P < 0.00001.

Intriguingly, while the non-hotspot T189M variant showed a cytoplasmic localization similar to the wild type RPL3L (**Fig. 4a-b**), the other non-hotspot R161W variant, identified in the same affected individual as the hotspot G27D, also mislocalizes to the nucleus (**Fig. 4a-b**). However, unlike the hotspot variants and consistent with its lower predicted pathogenicity (**Fig. 1c**), R161W does not cause nuclear sequestration of 28S rRNA (**Fig. 4a/c)**, nor does it lead to a ribosome biogenesis defect (**Extended Data Fig. 2a**) or cell toxicity (**Extended Data Fig. 2b**).

To gain biochemical insights into the toxicity associated with the hotspot variants, we profiled the interactomes of RPL3L variants by using immunoprecipitation against HA-tag followed by mass spectrometry in triplicates. Given the exclusive nuclear localization of the hotspot variants, we used nuclear lysate as input for all samples. Interestingly, principal component analysis (PCA) of the proteomics data separates the two hotspot variants from the rest in the first principal component (PC1), suggesting dramatic rewiring of the interactome for hotspot variants (**Fig. 4d**). The second principal component (PC2) separates the three nuclear variants (G27D, D308N, and R161W) from the two cytoplasmic variants (T189M and WT). Interestingly, the top two proteins driving the separation of hotspot variants from other variants both exhibit increased affinity for the hotspot variants, supporting the concept of toxic gain-of-function effects (**Fig. 4e**). These two proteins, GRWD1 and C7ORF50, although less studied in humans, have yeast homologs (Rrb1 and Rbp95, respectively) that directly interact with yeast Rpl3 during 60S ribosome biogenesis. GRWD1 serves as the dedicated chaperone for RPL3 (and likely RPL3L as well), binding RPL3 co-translationally to prevent its aggregation and degradation, ensuring its proper delivery to the assembly site on nucleolar pre-60S particles^14–16^. The other protein, C7ORF50, interacts with RPL3 both genetically and physically, binding adjacent to RPL3 on the 28S rRNA during early pre-60S ribosome biogenesis^49–51^. Several other 60S biogenesis factors, including RRP15^52^, EBNA1BP2^53^, DDX24^54^, NOP56^50^, and MRTO4^55^ also exhibited enrichment in the nuclear pulldown of RPL3L variants (**Extended Data Fig. 3**). The sequestration of these essential 60S ribosome biogenesis factors by hotspot variants likely underlies the 60S biogenesis defect and contributes to the toxic gain-of-function effects associated with these variants.

In summary, our subcellular localization analysis and nuclear protein interactome profiling using isogenic cell lines reveal that the hotspot variants D308N and G27D mislocalize to the nucleus, leading to nuclear retention of 28S rRNA. This toxic effect is likely driven by their enhanced binding to the dedicated RPL3/RPL3L chaperone GRWD1 and 60S ribosome biogenesis factors, including C7ORF50.

### Non-hotspot variants drive post-transcriptional *RPL3* compensatory up-regulation

It is intriguing that the mislocalization of the non-hotspot variant R161W to the nucleus (**Fig. 4a-b**) did not cause ribosome biogenesis defects (**Fig. 4c** and **Extended Data Fig. 2a**) or cell toxicity (**Extended Data Fig. 2b**). Western blot analysis of the polysome fractions confirmed the absence of R161W from these ribosomes (**Extended Data Fig. 4**). To investigate how ribosomes can still function in the absence of cytoplasmic RPL3L, we examined the expression of RPL3, which was expected to be silenced by an shRNA induced alongside the R161W variant upon dox treatment (**Fig. 3a**). Unexpectedly, while *RPL3* remained silenced in cells expressing most *RPL3L* variants, its expression was almost fully restored in cells expressing the R161W variant, and to a lesser extent in those expressing the T189M variant, at both the mRNA (**Fig. 5a**) and protein (**Fig. 5b**) levels. Importantly, there was no significant change in the expression of the *RPL3*-targeting shRNA (**Extended Data Fig. 5**), ruling out the possibility that *RPL3* recovery was due to loss of shRNA-mediated knockdown in cells expressing R161W or T189M. These results suggest that, similar to *Rpl3l*-KO mice, the expression of the non-hotspot *RPL3L* variants R161W and T189M leads to compensatory up-regulation of *RPL3*, suggesting that these variants are functionally equivalent to a knockout of *RPL3L*. Supporting a model in which compound heterozygotes carry a toxic hotspot variant alongside a loss-of-function variant, frameshifting or splice variants were identified in four of the eight families with hotspot variants (**Fig. 1a**).

**Figure 5.**
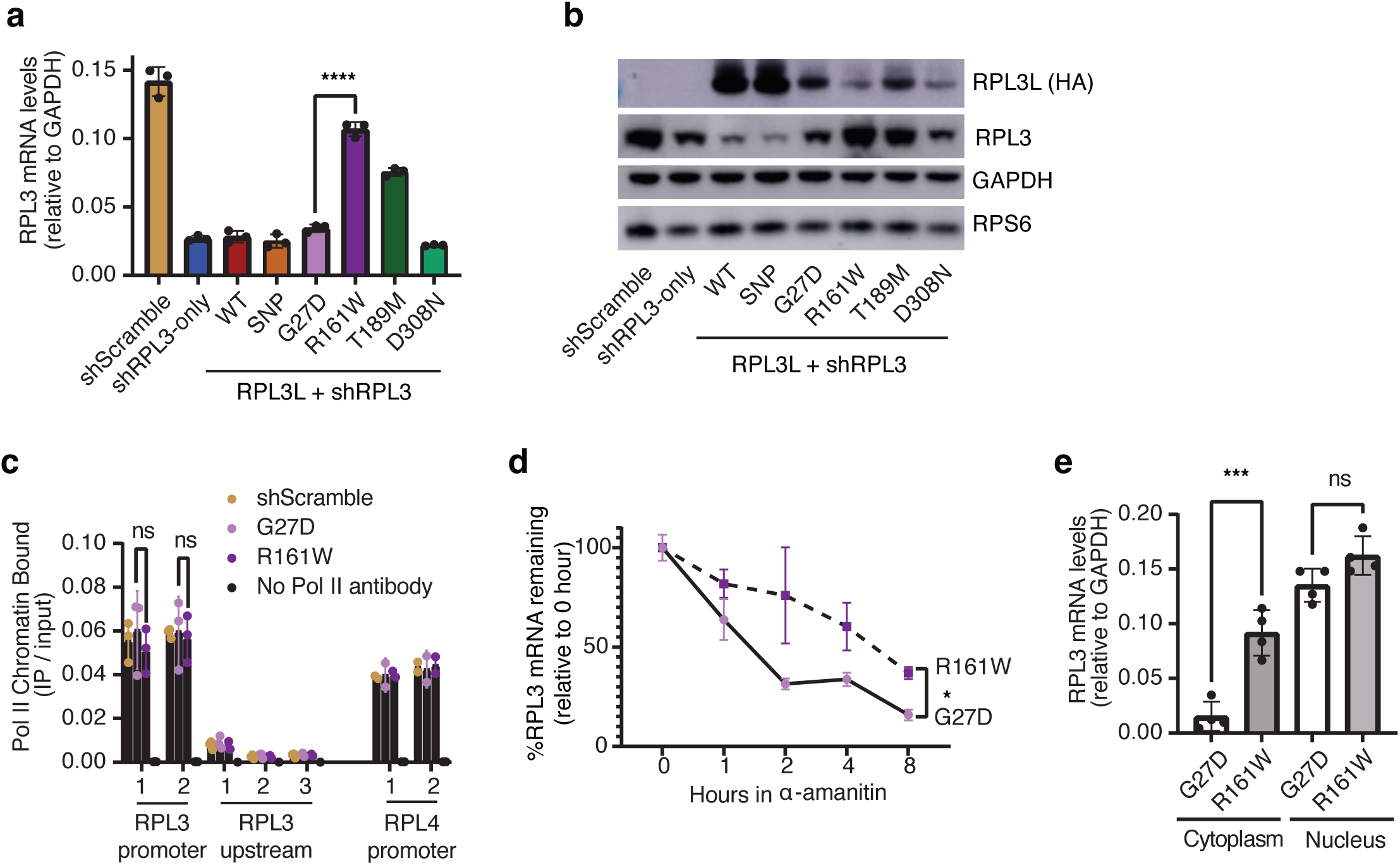
Non-hotspot variants drive post-transcriptional *RPL3* compensatory up-regulation. **a**, Compensatory increase of *RPL3* mRNA in R161W cells. DCM-associated *RPL3L* variants were expressed for 120 hours in doxycycline and total RNA was extracted and analyzed for *RPL3* mRNA levels (N = 3 independent experiments). **b**, Western blotting shows compensatory increase of RPL3 protein levels in R161W cells. Cells were treated as in (a) and whole-cell lysates were immunoblotted with indicated antibodies (N = 3 independent experiments). **c**, No difference in transcription activity in *RPL3* promoter assayed by ChIP-qPCR. Multiple primer pairs were used for each region. An intergenic region upstream of *RPL3L* was used as a negative control of non-transcribed region (N = 3). **d**, *RPL3* mRNA half-life is enhanced in R161W cells. *RPL3L*-expressing variants were incubated with 50μg/ml RNA Polymerase II inhibitor α-amanitin for indicated timepoints prior to analysis for *RPL3* mRNA relative to GAPDH, normalized to 0 hour (N = 3). **e**, Compensatory *RPL3* mRNA increase occurs entirely in the cytoplasm in R161W cells. RNA underwent subcellular fractionation for amplification of *RPL3* levels relative to GAPDH (N = 4). **a,c,d,e**, Data are mean ± s.d. ns, not significant; * P < 0.01; *** P < 0.0001; **** P < 0.00001.

To elucidate the mechanism behind this *RPL3* compensation, we first examined the transcriptional activity at the *RPL3* promoter using chromatin immunoprecipitation (ChIP) against RNA polymerase II (Pol II). No significant differences in transcriptional activity were observed between cells expressing G27D and R161W (**Fig. 5c**). Additionally, qPCR measurements of nascent transcription from intronic RNA showed no significant differences (**Extended Data Fig. 6**), ruling out transcriptional up-regulation as the mechanism of *RPL3* compensation.

We then assessed the stability of *RPL3* mRNA by inhibiting global transcription with the RNA Pol II inhibitor α-amanitin and measuring the decay of *RPL3* mRNA over time. Compared to cells expressing the G27D variant, *RPL3* mRNA was significantly more stable in cells expressing the R161W variant (**Fig. 5d**, P < 0.05). Consistent with a post-transcriptional mechanism, the increase in steady-state *RPL3* level was primarily due to increased cytoplasmic *RPL3*, with no significant change in the nuclear fraction (**Fig. 5e**).

In summary, unlike the two hotspot variants, G27D and D308N, which exhibit a gain-of-toxic function, the non-hotspot variants R161W and T189M resemble a knockout effect, promoting compensatory upregulation of *RPL3* by stabilizing its mRNA.

### Impaired protein synthesis in engineered compound heterozygous cells

*RPL3L*-linked DCM patients often carry a combination of a hotspot variant and a non-hotspot variant, such as the D308N/T189M and G27D/R161W pairs we have characterized. In our analysis of explanted heart tissue from patients carrying the D308N/T189M variants, both variants were detected at the RNA (**Fig. 2d**) and protein levels (**Fig. 2f**). Given the opposing effects of the hotspot and non-hotspot variants on ribosome biogenesis and cell growth (**Fig. 3-5**), we sought to explore the impact of these variants when both are present in the same cell, as is common in most DCM patients.

To address this, we introduced the G27D variant into cells expressing the R161W variant, creating a model of compound heterozygous cells that mirror those found in DCM patients. We chose R161W over T189M because R161W triggers a stronger compensatory up-regulation of *RPL3* (**Fig. 5a-b**). Using a puromycin incorporation assay to measure nascent protein synthesis, we observed a significant reduction in protein synthesis rates in the compound heterozygous cells compared to those expressing R161W alone, though the rate was still higher than in cells expressing G27D alone (**Fig. 6a-b**). This reduced protein synthesis rate aligns with the observed detrimental effects of compound heterozygous *RPL3L* variants in humans.

**Figure 6.**
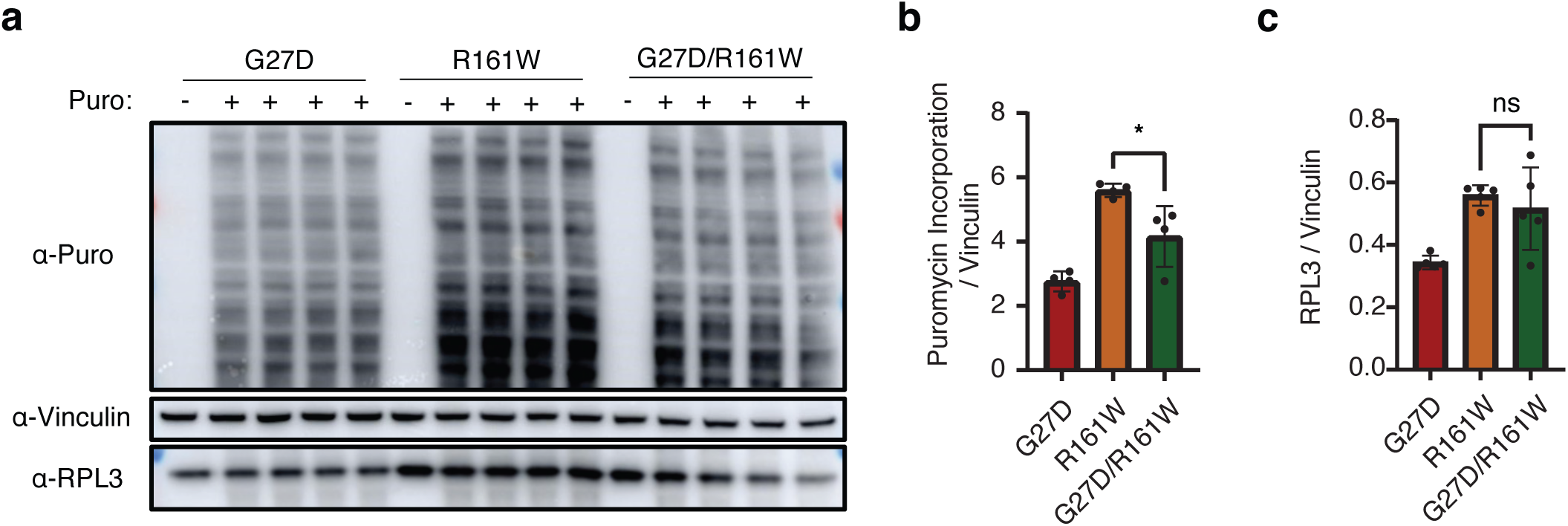
Impaired protein synthesis in engineered compound heterozygous cells. **a**, Cells stably integrated with G27D, R161W, or both G27D and R161W were induced with dox for 144hrs. Puromycin was added to 1uM final concentration for 30 minutes before cells were harvested for Western blotting (N = 4). **b**, Quantification of puromycin incorporation. **c**, Quantification of RPL3 protein abundance. Data are mean ± s.d. ns, not significant; * P < 0.01.

Interestingly, *RPL3* compensation remained unaffected in the compound heterozygous cells compared to cells expressing R161W alone (**Fig. 6c**). This suggests that the compensatory up-regulation of *RPL3* is not sufficient to overcome the toxic effects of the G27D variant.

## Discussion

In this study, we uncovered a unifying mechanism linking a large number of variants in muscle-specific ribosomes to severe neonatal heart failure. Remarkably, affected families often carry one of two toxic gain-of-function variants alongside a unique putative loss-of-function variant. The loss-of-function variants include frameshift-induced truncated variants and missense variants distributed across the protein, consistent with the overall high evolutionary conservation of RPL3L. This unexpected pathogenetic mechanism—autosomal recessive mutations exerting gain-of-toxic function other than merely loss-of-function effects—underlie the severe heart defects in humans and may also explain the lack of phenotype in *RPL3L* KO mice and humans.

Our results show that the disease is primarily driven by two hotspot variants exhibiting toxic gain-of-function effects on ribosome biogenesis. In rare diseases, hotspot variants are frequently associated with toxic effects or dominant-negative traits^56–58^. Despite their rarity in the general population, the missense variants G27D and D308N/V are present in eight of ten unrelated families affected by DCM (**Fig. 1a**). Mechanistically, these RPL3L hotspot variants mislocalize to the nucleus (**Fig. 4a-b**) and drastically rewire the network of 60S ribosome biogenesis factors (**Fig. 4d-e**). Supporting a toxic gain-of-function model, both hotspot variants exhibit enhanced binding to the dedicated RPL3/RPL3L chaperone GRWD1 and several early pre-60S biogenesis factors (**Fig. 4d-e**). This aberrant interaction leads to sequestration of 28S rRNA in the nucleus (**Fig. 4a/c**), a defect absent in cells expressing other variants or lacking RPL3L expression entirely, underscoring the specific toxicity of the hotspot variants beyond simple loss-of-function. Expression of the hotspot variants results in a marked depletion of ribosomes (**Fig. 3e**) and a severe cell growth defect (**Fig. 3f**), effects that extend beyond the consequences of ribosome loss alone. Future studies, especially *in vivo* studies will be needed to elucidate how these molecular and cellular defects lead to organ level failure.

Autosomal recessive mutations in Mendelian disorders are typically considered loss-of-function mutations^59^. Rarely do they cause disease via gain-of-toxicity or dominant-negative mechanisms. The only prior example known is the A673V missense mutation in the β-amyloid (Aβ) precursor protein (APP), associated with familial Alzheimer’s disease^33^. In this case, the mutant in the homozygous state enhances Aβ peptide production, promoting the formation of amyloid fibrils, the primary trigger of Alzheimer’s disease. In heterozygous individuals, however, the coexistence of wild-type and mutant Aβ peptides inhibits amyloidogenesis relative to either mutant or wild-type Aβ peptide alone. This inhibition is likely due to conformational incompatibility between the wild-type and mutant Aβ peptides. For *RPL3L*-linked heart failure, there are several possibilities for why heterozygous members of families carrying the recurrent and toxic variant (e.g., G27D and D308N/V) have not developed severe disease. The presence of the wild type RPL3L protein will likely reduce the toxicity and result in incomplete penetrance or much delayed onset of the disease. Given that yeast Rpl3 aggregates when over-expressed^14^, one plausible mitigation mechanism is that these RPL3L hotspot variants promote aggregation similar to the A673V Aβ variant, with wild-type RPL3L in heterozygous cells preventing such aggregation and mitigating the pathogenic effects.

Compensation by the paralog RPL3, triggered by RPL3L loss-of-function, likely explains why severe disease arises only from a combined gain-of-toxicity and loss-of-function mechanism, rather than loss-of-function alone. Given that 80% Mendelian disease genes have duplicated paralogs^60^, similar combinatorial mechanisms may be more widespread. Genetic compensation by paralogs is well-documented, although the underlying molecular mechanisms often remain unclear^61–63^. The increased expression of *Rpl3* observed in *Rpl3l* KO mice has been proposed to explain the absence of severe cardiac phenotypes in these animals, but the molecular basis of this compensation is not yet understood. In this study, we demonstrate that non-hotspot missense mutations, such as R161W, also up-regulate *RPL3* expression in human cells. This up-regulation occurs through stabilization of cytoplasmic *RPL3* mRNA without transcriptional activation. However, the *RPL3* compensation induced by the R161W variant does not fully counteract the toxic effects of the G27D variant when co-expressed in human cells. These findings suggest that enhancing *RPL3* compensation alone may be insufficient as a therapeutic strategy for patients carrying toxic *RPL3L* mutations.

We observed no significant *RPL3* compensation in failing hearts explanted from human DCM patients carrying the compound heterozygous RPL3L variants, D308N and T189M. This finding stands in stark contrast to the robust up-regulation of *Rpl3* observed in *Rpl3l* KO mice. The absence of *RPL3* compensation in humans may contribute to the disease pathology and potentially explain the phenotypic differences between humans and mice. Alternatively, it could result from muscle remodeling and other pathological changes in failing hearts, given that the RPL3/RPL3L switch is regulated by muscle hypertrophy. Supporting the latter hypothesis that the absence of *RPL3* compensation is a consequence rather than a cause of the disease, *RPL3* is down-regulated in non-*RPL3L*-linked DCM^64^. Consistent with this model, *RPL3* compensation was robustly induced *in vitro* in AC16 cells co-expressing same variants (D308N/T189M) found in patient cells. These findings suggest that *RPL3* compensation may occur during the early stages of the disease but diminishes as the condition progresses. The disease can progress even in the presence of *RPL3* compensation, as shown *in vitro* that the D308N variant causes protein synthesis defect even in the presence of *RPL3* compensation (**Fig. 6**). As the disease advances, the loss of *RPL3* compensation could result from the complex remodeling processes in failing hearts. Future studies, including mouse knock-in models that faithfully recapitulate human disease phenotypes, will be critical for testing these models and uncovering the underlying mechanisms.

In conclusion, our systematic genetic and functional analyses shed light on the pathogenesis of heart failure caused by mutations in muscle-specific ribosomes, revealing a novel mechanism driven by the gain-of-toxic effects of recessive mutations. These findings provide a foundation for exploring other RPL3L variants and developing targeted therapies. Notably, our results suggest that gain-of-toxicity mechanisms may be more prevalent in autosomal recessive diseases than previously recognized. Furthermore, the interplay between gain-of-toxicity and loss-of-function mechanisms may represent a common feature in diseases involving genes with paralogs. Finally, our study highlights unexpected tissue-specific pathology in otherwise universal cellular processes and underscores the importance of unbiased approaches in identifying disease candidates for accurate diagnosis.

## Acknowledgments

We thank members of the Wu lab and the Cardiometabolic Genomics Program for discussion and comments on the manuscript. This work is supported by NIH/NHLBI grant 1R01HL171664-01 and Pershing Square Foundation MIND Prize (X.W.) and a postdoctoral fellowship from American Heart Association (M.R.M). This research was funded in part through the NIH/NCI Cancer Center Support Grant P30CA013696 and used the Genomics and High Throughput Screening Shared Resource and CCTI Flow Cytometry Core. The CCTI Flow Cytometry Core is supported in part by the Office of the Director, National Institutes of Health under awards S10OD030282 and S10OD020056.

## Author contributions

X.W. and M.R.M. conceived the functional study and wrote the manuscript with input from all authors. M.R.M. performed the majority of experiments. M.G. coordinated case reports sample collection and provided critical feedback. T.M.L. and J.M.F. identified and characterized the D308N/T340Nfs*26 case. M.V.P., A.B., and P.J. identified and characterized the D308N/R58Q case. A.L.R., R.C.A., and D.N. identified and characterized the D308N/c.1167+1G>A case. R.K.S. generated the proteomics data. Y.Y. assisted in imaging. W.K.C. contributed the D308N/T189M tissue sample. W.K.C., M.P.R., and F.Y. edited the manuscript and provided critical feedback.

## Data availability

RNA-seq data has been deposited in GEO. It will be released after publication.

## Code availability

Data analyses were performed using publicly available software.

## Materials and Methods

### Cell Culture, Drugs and Transfection

AC16 cells were cultured at 37°C, 5% CO_2_ in low-glucose DMEM (Fisher; 11-885-084) supplemented with 10% FBS. HEK293T cells were cultured in DMEM with 10% FBS. For immunofluorescence assays, 2 x 10^5^ cells were plated in 6 well plates and incubated for 24 hours. For stable cell generation, EZ-Tet-RPL3-shRNA was co-transfected with CMV-dR8.91 and MD2.G plasmids for 72 hours in HEK293T cells to produce virus. The media was centrifuged for 5 minutes at RT, filtered using 0.45µm Syringe Filter Unit (Fisher), and 200µl viral media was added to 2 x 10^5^ AC16 cells for 48 hours prior to selection using 200µg/ml Hygromycin B until control cells (non-transduced) were dead. For stable generation of pLenti-RPL3L, procedure was followed as above except transduction was carried out on EZ-Tet-RPL3-shRNA/AC16 cells and selection was carried out by sorting for BV421-positive cells. For mRNA stability assay, AC16 cells were treated with 50mg/ml a-amanitin for the indicated time points.

### Plasmids

To generate plasmid expressing shRNA against RPL3, a reported high efficiency shRNA sequence for RPL3 was identified (Sigma) and ∼70nt single stranded oligos were ordered from IDT and annealed by heating to 95°C and cooled at a rate of 5°C min^-1^. Double-stranded oligos had two 5’ overhangs corresponding to NheI and EcoRI sites, respectively. dsDNA oligos were ligated to EZ-Tet-pLKO-Hygro (Addgene; 85972) using Quick Ligation Kit (NEB), producing Tet-inducible RPL3 shRNA vector. The pLenti-RPL3L plasmid was derived from pLentiRNACRISPR_007 (Addgene; 138149). A gene fragment containing the wild-type human RPL3L sequence with a C-terminal HA tag was delivered on pUC57 (Genewiz). PCR products for wild-type RPL3L-HA derived from pUC57, the pLentiRNACRISPR_007 backbone, and EBFP (unpublished in-house plasmid) with overlapping ends were assembled via Gibson cloning using NEBuilder HiFi DNA Assembly Master Mix (NEB), producing Dox-inducible RPL3L and constitutive EBFP expression. To obtain RPL3L mutation plasmids, primers (Supplementary Table) were designed against pUC57-RPL3L-HA with single nucleotide substitutions in the 5’ region of the forward primer sequence. Mutagenesis was performed using Site Directed Mutagenesis Kit (NEB) as per the manufacturer’s instructions. pLenti-RPL3L mutations were derived similarly to the wild-type as above.

### Immunofluorescence and FISH

15,000 cells were seeded on Lab-Tek II (Thermo) chamber slides and grown overnight. Cells were washed once with PBS and fixed in 300µl PFA at room temperature for 15 minutes. Fixed wells were washed 3x with PBS for 3 minutes, permeabilized in 0.5% Triton X-100 in PBS for 15 minutes, washed 3x, and incubated in 3% BSA in PBS for 1 hour to block. Cells were then incubated in 3% BSA containing primary antibody overnight at 4°C, washed 3x in PBS, then incubated for 1 hour at room temperature in secondary antibody protected from light. After washing 3x, cells were incubated in 1:1000 DAPI solution for 5 minutes, washed 3x, then imaged using a Nikon Eclipse Ti Series confocal microscope equipped with 60x lens performed at Columbia University’s Confocal and Specialized Microscopy Core. RPL3L was detected using anti-HA (Thermo; 2-2.2.14). Secondary staining was accomplished using goat anti-rabbit IgG (H+L) Alexa Fluor 488 (Invitrogen) and goat anti-mouse IgG (H+L) Alexa Fluor 594 (Invitrogen). For rRNA FISH, cells were fixed as above but permeabilized with 70% ethanol for at least 1 hr. Custom designed 28S rRNA FISH probes (Supplementary Table) were designed conjugated to CAL Fluor® Red 590 dye (Stellaris) and incubated with fixed samples for 16 hours in the dark. After washing using wash buffers A + B (Stellaris), samples were incubated with primary anti-HA antibody as above.

### Polysome fractionation

Sucrose gradient density ultracentrifugation was used to separate the monosomal and polysomal ribosomal fractions. For human cell experiments, AC16 cells were grown in 15cm dish were lysed by adding 800µl of ice-cold lysis buffer (20mM HEPES, 150mM KCl, 10mM MgCl_2_), plus 0.5% NP-40, 1x EDTA-free Protease Inhibitor Cocktail (PIC; Roche)), 2.5mM DTT, 25 U ml^−1^ Turbo DNase I (Invitrogen)), 10U SuperaseIN (Thermo), 2mM Vanadyl Ribonucleoside Complex (NEB) and 100ug/ml CHX (Sigma) directly to the plate and scraping. Lysed material was transferred to 1.5ml centrifuge tube then triturated 10x with 23-gauge syringe. Samples were incubated in 4°C for 30 minutes with rotation to complete lysis, then centrifuged at 14,000g for 7 minutes. Supernatant was transferred to new tube and 700ul of lysate was loaded onto a 10-50% sucrose gradient, before ultracentrifugation at 38,000rpm for 2 hours. The sucrose gradient was made by combining 10% and 50% sucrose solutions in lysis buffer, plus 2.5mM DTT, 160U SuperaseIN, and 100µg/ml CHX. Both gradient formation and fractionation was performed on a Biocomp Gradient Profiler (Biocomp). For protein extraction post-fractionation, 1 volume of TCA to 4 volumes of protein sample was added, incubated for 10 minutes on ice, then span at 14,000rpm for 5 minutes at 4°C. Pellet was washed twice with ice-cold acetone and dried at 95°C for 5 minutes. For SDS-PAGE, 4x Sample reducing agent (GenScript) diluted 1:1 in RIPA was added to dry pellet and heated for an additional 5 minutes at 95°C.

### Western blotting

Western blot analysis was performed after SDS-PAGE using standard protocols. The antibodies used were anti-HA (Thermo; 2-2.2.14), anti-RPL3 (Sigma; 3365), anti-RPL28 (Invitrogen; 62192), anti-RPS3 (Thermo; 2G7H4), anti-GAPDH (Invitrogen; PA1988), and anti-Vinculin (CST; E1E9V).

### RNA extraction/analysis and qPCR

Total RNA was isolated using NucleoSpin RNA Plus kit (Macharey-Nagel) including use of gDNA removal column. For rRNA analysis, 500ng RNA was ran on 2100 Bioanalyzer (Agilent) at the Columbia University’s Molecular Pathology Core. For qPCR, cDNA synthesis was performed using 500ng RNA with the SuperScript IV Reverse Transcriptase (Invitrogen) as per the manufacturers protocol with oligo(dT) primers. Real-time PCR was carried out using PowerUp SYBR Green Master Mix using 1:3 diluted cDNA. For qRT-PCR, triplicates were performed with 0.5µM primer and 1µl cDNA per reaction in the QuantStudio 7 Flex Real-Time PCR System. Ct values obtained were plotted using GAPDH mRNA as loading control. Quantifications were performed as per the MIQE guidelines. Statistical tests were performed with Prism 9 and two-sided Student’s t-test were used to calculate the *P* values. Data are expressed as mean ± s.d. for the number of replicates indicated, with *P* < 0.01 considered significant. All primer sequences are displayed in Supplementary Table.

### Nuclear/Cytoplasmic RNA Extraction Protocol

Cells were washed with ice-cold PBS, scraped into 1.7ml tubes, and pelleted at 500 g for 5 minutes at 4°C. Cytoplasmic RNA was extracted by lysing cells in Cytoplasmic Lysis Buffer (40 mM HEPES-NaOH pH 7.5, 160 mM KCl, 10 mM MgCl₂, 0.5% NP-40, 0.5% glycerol, and SuperaseIn) for 30 minutes at 4°C with rotation, followed by centrifugation at 2000 g for 2 minutes to separate the cytoplasmic supernatant. Nuclear pellets were washed in Cytoplasmic Lysis Buffer (without detergent), centrifuged at 500 g for 5 minutes, and subjected to a wash with Nuclear Lysis Buffer (10 mM Tris-HCl pH 8.0, 1.5 mM KCl, 2.5 mM MgCl₂, 5% glycerol, 0.5% Triton X-100, 0.5% deoxycholate, and 20U SuperaseIN) for 15 minutes at 4°C. Final washes were performed with Cytoplasmic Lysis Buffer (without detergent), and nuclei were pelleted at 2000 g for 2 minutes. Both cytoplasmic and nuclear RNA were extracted as above.

### Co-immunoprecipitation

To prepare magnetic bead-antibody complex, 50µl of Dynabeads Protein G (Invitrogen) was washed 3x with lysis buffer then incubated with 5µl anti-HA (Thermo; 2-2.2.14) in 400µl lysis buffer for 1 hour at RT with constant rotation. Magnetic bead-antibody complex was captured using Magnetic Separation Rack (NEB), washed 2x with cytoplasmic lysis buffer (20 mM Tris-HCl, pH 7.5, 150 mM KCl, 10% glycerol and 0.1% NP-40) then 3x with lysis buffer. 2x 15cm plates at 80% confluency were grown, cells were washed twice in ice-cold PBS and 400µl of ice-cold lysis buffer was dripped onto the plate and scraped, incubated in microcentrifuged tubes after triturating 10x with 23-gauge syringe, and centrifuged at 14,000g for 7 minutes. The supernatant was added to bead-antibody complex and incubated with rotation at 4°C overnight. Next day, bead-antibody-antigen complex was washed 3x for 10 minutes at 4°C with ice-cold lysis buffer. Protein was eluted by incubation with 40µl pH 3 glycine and heating at 70°C with shaking for 10 minutes, then neutralized with 1/10 volume pH 8.0 Tris.

### Proteomics

Explanted frozen patient tissue was homogenized with pestle and mortar under liquid nitrogen and lysed in RIPA for mass spectrometric analysis by the Proteomics core facility at Columbia. For the global quantitative proteomic analysis, diaPASEF-based proteomics^65^ was employed. Briefly, frozen patient tissue samples were lysed in freshly prepared SDC lysis buffer (1% sodium deoxycholate and 100 mM Tris-HCl, pH 8.5) and heated at 60°C with shaking at 1200 rpm for 15 minutes^66^. Protein reduction and alkylation of cysteine residues were carried out using 10 mM tris(2-carboxyethyl)phosphine (TCEP) and 40 mM chloroacetamide (CAA) at 45°C for 15 minutes, with shaking at 1200 rpm. The samples were then sonicated in a water bath and allowed to cool to room temperature. Proteins were digested overnight with a Lys-C/trypsin mix at a 1:50 ratio (µg enzyme to µg protein) at 37°C with shaking at 1400 rpm. The resulting peptides were acidified with 1% trifluoroacetic acid (TFA), vortexed, and subjected to StageTip cleanup using SDB-RPS^66^. Peptides were then dried in a speed vacuum concentrator. Dried peptides were resuspended in 10 µL of LC buffer (3% acetonitrile, 0.1% formic acid), and their concentrations were determined using a NanoDrop spectrophotometer. For diaPASEF analysis on the timsTOF Pro instrument, 200 ng of peptides from each sample were used. For liquid chromatography with tandem mass spectrometry (LC-MS/MS), peptides were separated within 87 min at a flow rate of 300 nl/min on a reversed-phase C18 column with an integrated CaptiveSpray Emitter (25 cm x 75µm, 1.6 µm, IonOpticks). Mobile phases A and B were with 0.1% formic acid in water and 0.1% formic acid in ACN. The fraction of B was linearly increased from 2 to 23% within 70 min, followed by an increase to 35% within 10 min and a further increase to 80% before re-equilibration. The timsTOF Pro was operated in diaPASEF mode and data was acquired at defined 32 × 50 Th isolation windows from m/z 400 to 1,200. To adapt the MS1 cycle time in diaPASEF, set the repetitions to 2 in the 16-scan diaPASEF scheme. The collision energy was ramped linearly as a function of the mobility from 59 eV at 1/K0=1.6 Vs cm^-2^ to 20 eV at 1/K0=0.6 Vs cm^-2^. The acquired diaPASEF raw files were searched using the UniProtKB/Swiss-Prot mouse database was performed using library-free workflow in the DIA-NN^67^ search engine, employing the default settings of the library-free search algorithm with match-between-runs (MBR) enabled. A maximum of 1 trypsin missed cleavage was allowed and the maximum variable modification was set to 1. Carbamidomethylation was set as the fixed modification, whereas protein N-terminal methionine excision, methionine oxidation, and N-terminal acetylation were set as variable modifications. The peptide length range was set to 7–30 amino acids, precursor charge range 1–4, precursor m/z range 300–1800, and fragment ion m/z range 200–1800. The false discovery rates (FDRs) at the protein and peptide levels were set to 1%, and MS1 and MS2 mass tolerances were automatically determined by DIA-NN. Results obtained from DIA-NN were subjected to further statistical analyses. Custom fasta files containing mutant D308N peptides were analyzed on MaxQuant for peptide abundance relative to wildtype peptide abundance. For nuclear IP of cell lines, MaxQuant outputs were analyzed using Differential Expression of Proteins (DEP) package in R.

### Chromatin Immunoprecipitation (ChIP)

Cells were crosslinked with 1% formaldehyde in the culture medium for 15 minutes at room temperature, followed by quenching with 125 mM glycine for 5 minutes. Cells were pelleted and washed. The pellet was resuspended in RIPA buffer with protease inhibitor cocktail (Roche), incubated on ice, and sonicated to shear chromatin (∼1 kb fragments). The lysate was centrifuged at 12,000 x g for 10 minutes at 4 °C, and the volume was increased with dilution buffer (165 mM NaCl, 0.01% SDS, 1.1% Triton X-100, 1.2 mM EDTA, 16.7 mM Tris–HCl, pH 8.0). Chromatin was incubated with 5 μg of anti-RNAPII (Santa Cruz; F-12) for overnight at 4 °C. Chromatin-Antibody mix was incubated with pre-washed Dynabeads and incubated at RT for 45 minutes. Beads were washed 5x in dilution buffer and eluted in genomic DNA extraction kit (Macharey-Nagel) lysis buffer, resuspended in 50 μL of water. PCR was performed using purified DNA and compared to input samples. Two-sided Student’s t-tests were carried out to calculate *P* value.

### Nuclear localization calculation

Nuclear localization was calculated as a ratio of the immunofluorescent co-localization signal of anti-HA with nuclear regions compared to anti-HA signal with cytoplasmic regions, determined as cellular regions lacking in DAPI signal. The same calculation was performed for 28S rRNA. All calculations were carried out with ImageJ. Cytoplasmic/Nuclear signal ratios for each cell were compiled and underwent one-way ANOVA with Dunnett correction for multiple comparisons across conditions.

### Tissue RNA-seq

Explanted frozen patient heart tissue was homogenized using pestle and mortar under liquid nitrogen. Homogenized tissue was submerged in TRIzol and extracted using Direct-zol kit (Zymo Research). RNA integrity was analyzed with Bioanalyzer electrophoresis. Samples underwent poly(A) RNA pulldown and libraries were constructed using Illumina TruSeq chemistry. Libraries were sequenced at Columbia Sulzberger Genome Center with NovaSeq 6000 (Illumina). RTA (Illumina) was used for base calling and bcl2fastq2 (version 2.19) for converting BCL to fastq format. Gene expression was quantified using kallisto with a customized transcriptome index (human MANE select). RNA-seq data for three age-matched control samples were downloaded from European Nucleotide Archive (ENA project PRJEB26969, sample ID: 5818sTS, 5828sTS, and 5836sTS) and analyzed similarly. DESeq2 was used for differential gene expression analysis.

## Figure legends

**Extended Data Figure 1.**
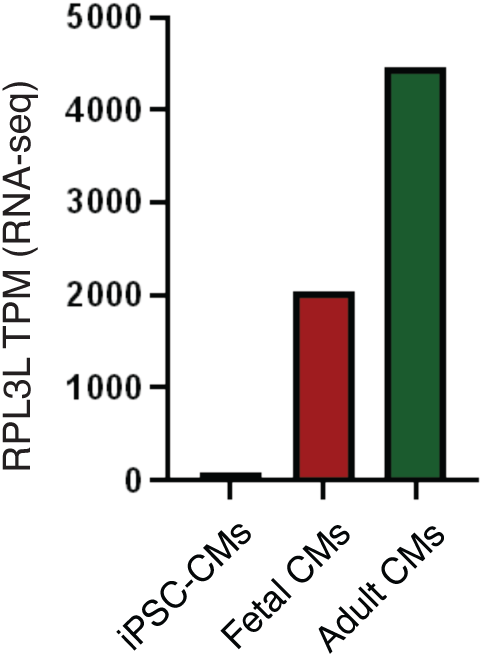
iPSC cardiomyocytes (iPSC-CMs) do not express *RPL3L*. Data was analyzed from RNA-seq performed by Pozo et al (2022) comparing iPSC-CMs to fetal and adult human heart.

**Extended Data Figure 2.**
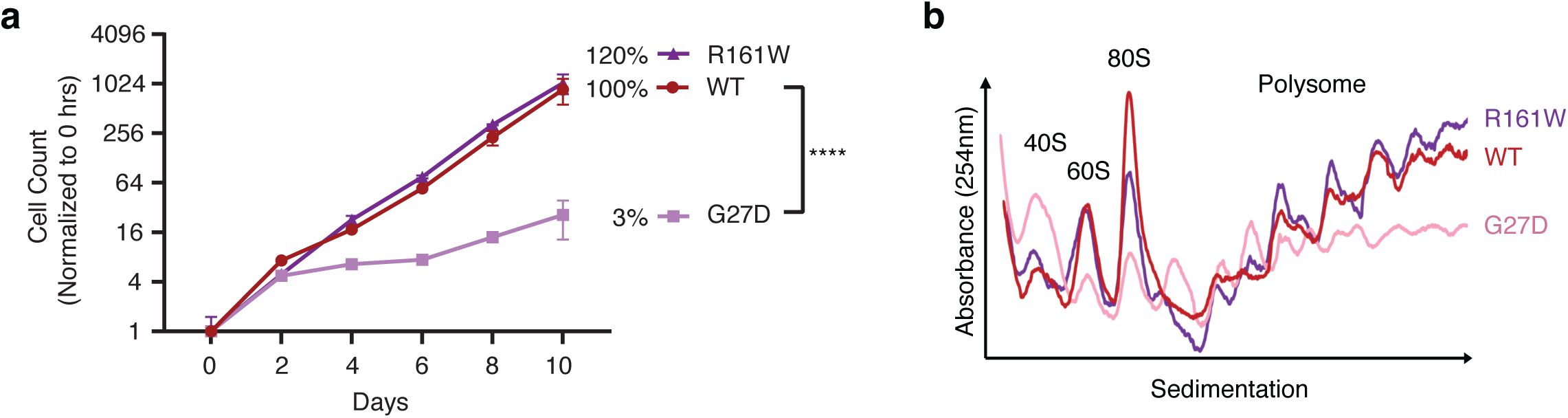
The hotspot G27D variant mirrors the effects of D308N. **a**, Cell growth defect for G27D and R161W, as in **Fig. 3f**. **b**, Polysome fraction for G27D and R161W, as in **Fig. 3e**. Data are mean ± s.d.

**Extended Data Figure 3.**
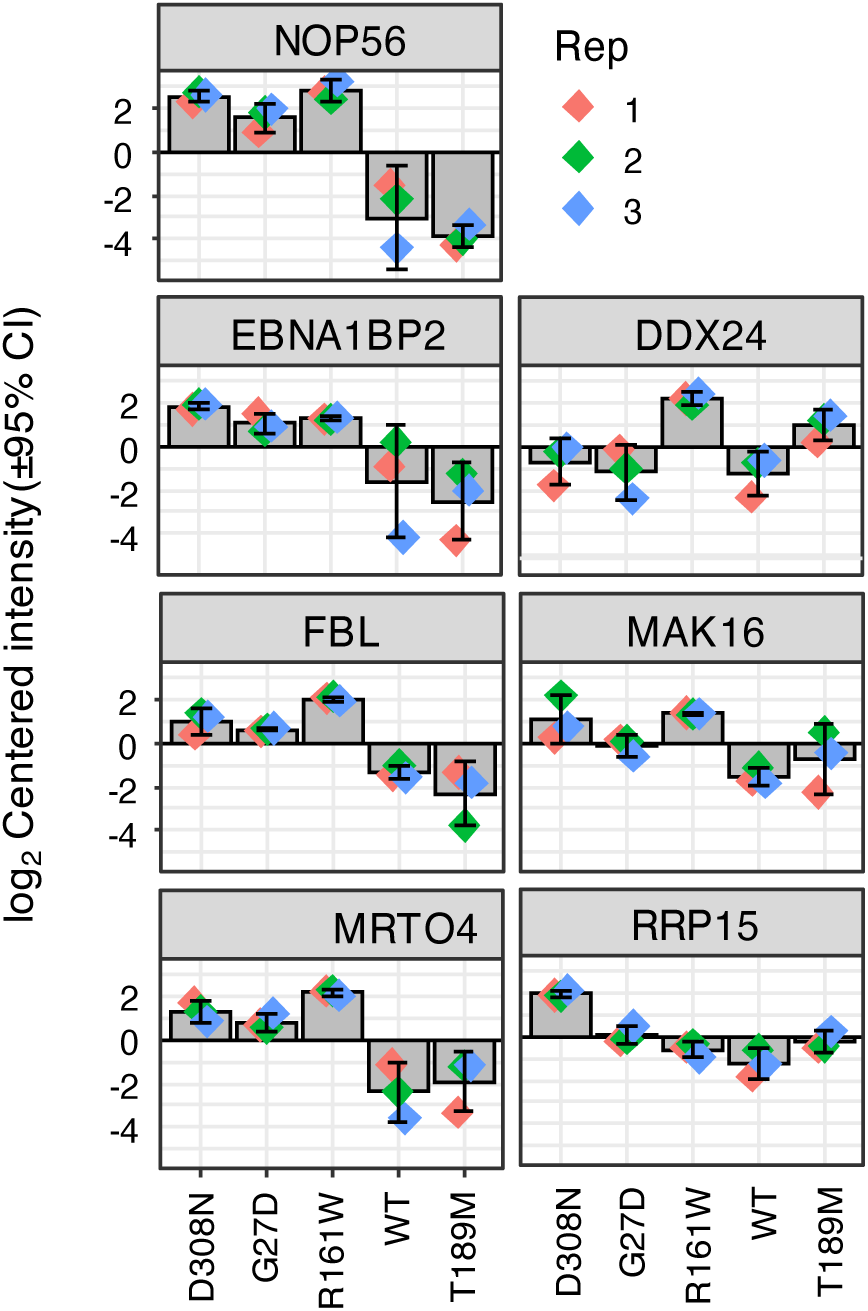
Enrichment of 60S ribosome biogenesis factors in proteins co-immunoprecipitated with nuclear RPL3L variants. Peptide intensities are normalized by the median of all samples and then log2-transformed. Error bars represent standard deviation.

**Extended Data Figure 4.**
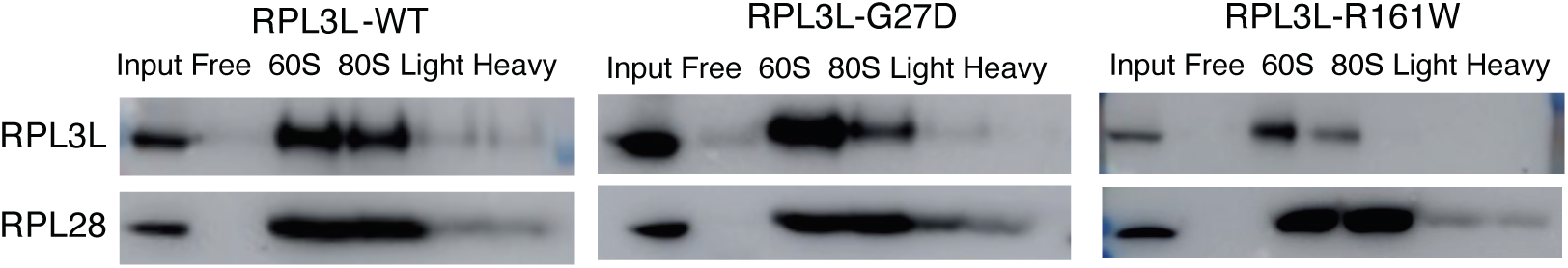
R161W does not incorporate into polysomes. Monosome and polysomal fractions from **Fig. 5** were isolated by TCA precipitation and immunoblotted for HA-tag. Input (pre-ultracentrifugation) samples were loaded as controls.

**Extended Data Figure 5.**
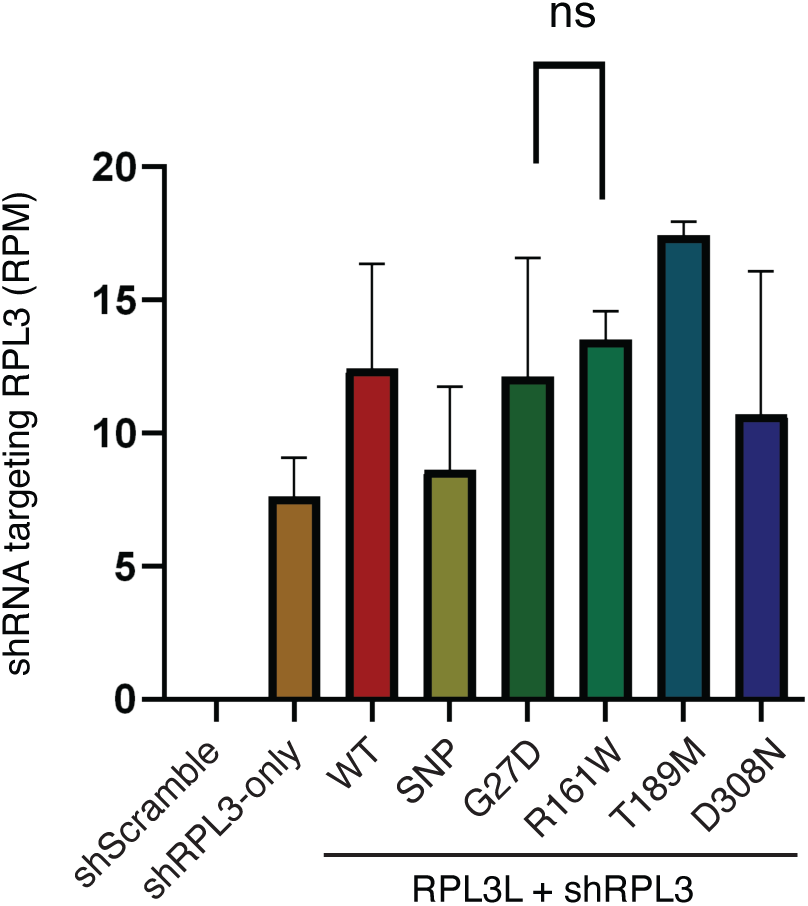
No change in the expression of the *RPL3*-targeting shRNA in *RPL3L*-expressing cells. A custom annotation was used to detect shRNA hairpin from RNA-seq data obtained from *RPL3L* variant-expressing cell lines. RPM, Reads per million (N = 3 biological replicates). Error bars represent standard deviation. ns, not significant.

**Extended Data Figure 6.**
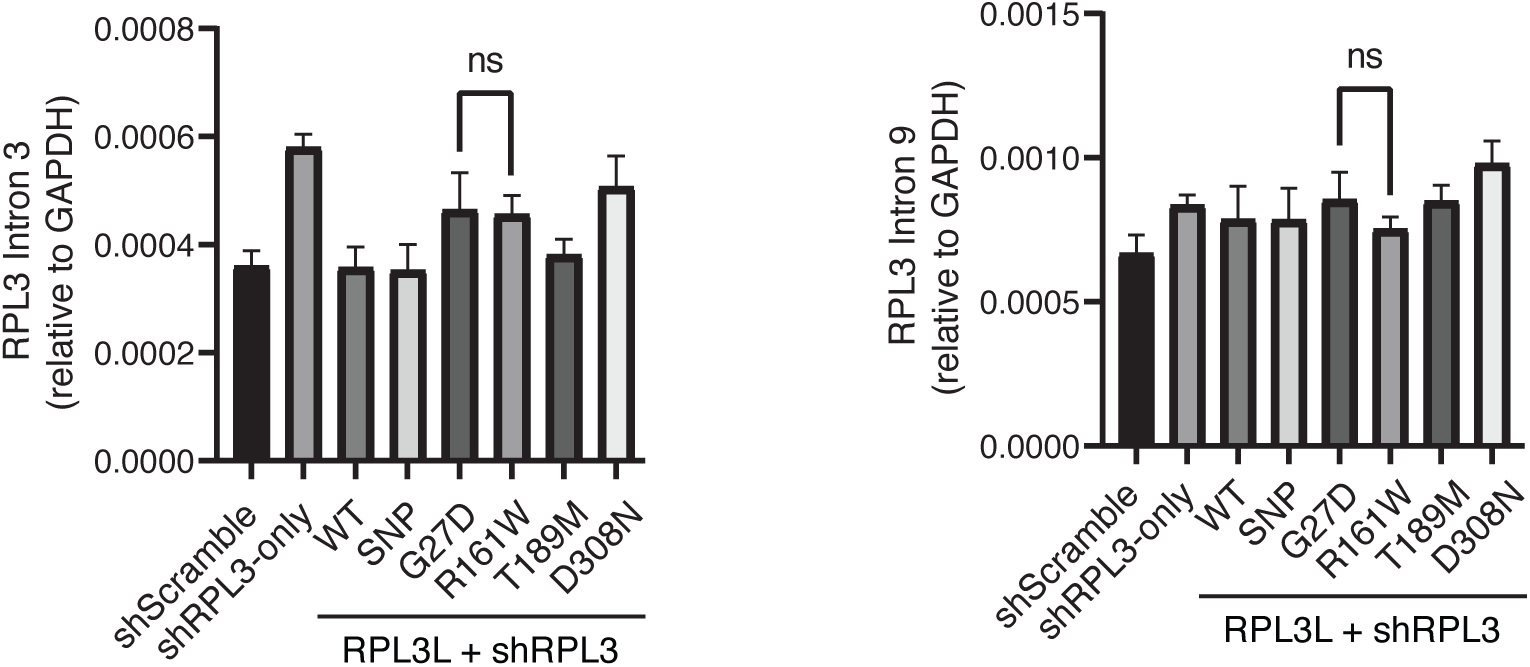
R161W does not induce higher *RPL3* intronic RNA as a proxy of transcription levels. Total RNA was isolated and cDNA was synthesized using random primers. Primers targeting intron 3 and intron 9 of *RPL3* were used for amplification (N = 3). Error bars represent standard deviation. ns, not significant.

