## Supplementary Information for "Pathogenetic mechanisms of muscle-specific ribosomes in dilated cardiomyopathy"

**Supplementary Note 1: case report**

**Case 1: R58Q/D308N**

The proband is a 3-month-old female, who presented with decreased oral intake, cough, and increased work of breathing over 5 weeks. She was born to a 32-year-old G1P1M0A0 mother at 36 weeks, twin gestation. On presentation, tachycardia and gallop were noted. Chest radiograph showed cardiomegaly with pulmonary congestion (Figure 1). EKG is shown in Figure 2. She was found to have congestive heart failure with reduced ejection fraction of 24%, with a severely dilated left heart. Her cardiac biomarkers were elevated with troponin of 0.11ng/ml and brain natriuretic peptide >3801 pg/ml. Metabolic labs including urine organic acids, carnitine panel, acylcarnitine panel, and newborn screen were normal. Infectious workup including CMV, Adenovirus, Coxsackievirus, Enterovirus, Hepatitis A, B, and C, Parvovirus, and Toxoplasmosis were negative. She was discharged home one month after hospitalization on an oral heart failure regimen including captopril, carvedilol, furosemide, aspirin, spironolactone, and digoxin.

Family history is unremarkable, with a 31-year-old mother of Cuban descent and a 33-year-old father of Colombian descent, with no consanguinity. Both parents are healthy with no medical conditions. Our proband has a twin brother who is healthy. Figure 3 describes the family pedigree. There is no known incidence of late-onset DCM in the family. Her twin brother has had several normal cardiac evaluations. Genetic testing is delayed per parental request. The family history is otherwise negative. Table 1 shows the clinical and molecular characteristics of the family.

The Cardiomyopathy panel for the proband was positive for two variants of uncertain significance HRAS- c.284A>G (p.Gln95Arg) and RYR2-c.7955T>C (p.Leu2652Pro), both maternally inherited from an asymptomatic mother. HRAS variant is now reclassified as Benign-mat inherited. TRIO Whole genome sequencing identified compound heterozygous variants of uncertain significance in the RPL3L gene - maternally inherited heterozygous missense variant, c.173G>A (p.Arg58Gln, R58Q) and paternally inherited heterozygous missense variant, c.922G>A (p.Asp308Asn, D308N). These variants in RPL3L were rare in the population databases.

At 19 months old, she remained on her heart failure regimen with addition of Entresto (Sacubitril/Valsartan) by her heart failure specialist. At 21 months of age, she had an acute decompensation requiring admission and inotropic support with Milrinone. Her echocardiogram showed progressively depressed systolic function. She was transferred to the transplant center and passed away from complications of heart failure prior to receiving a heart.

| Individual | **Age/ Gender** | **Cardiac phenotype** | **Management** | **Electro-cardiography** | **Echo-cardiography** | **Abnormal biomarkers upon admission** | **Genotype** | **Zygosity** |
|---|---|---|---|---|---|---|---|---|
| **Proband** | 3 month / Female | Dilated cardiomyopathy | Intravenous followed by oral heart failure therapy | Sinus Rhythm Normal axis and intervals<br>Possible right atrial enlargement Possible LVH | Severely dilated LA and LV<br>Severe MR LVEF 24% | Troponin 0.11ng/ml BNP >3801 pg/ml | c.173G>A<br>(p.Arg58Gln) and c.922G>A<br>(p.Asp308Asn) | Compound Heterozygous |
| **Mother** | 31 years / Female | Normal | Longitudinal clinical surveillance | | | | c.173G>A<br>(p.Arg58Gln) | Heterozygous |
| **Father** | 33 years / Male | Normal | Longitudinal clinical surveillance | | | | c.922G>A<br>(p.Asp308Asn) | Heterozygous |
| **Twin Brother** | 3 month / Male | Normal | Longitudinal clinical surveillance | | Normal | N/A | N/A | N/A |

**Table 1: Clinical and molecular characteristics of the family members (Case 1).**

Table legends: BNP, brain natriuretic peptide; LA, Left atrium; LV, left ventricle; LVEF, left ventricular ejection fraction; LVH, left ventricular hypertrophy; Hgb, hemoglobin; N/A, not applicable; VUS, a variant of uncertain significance.

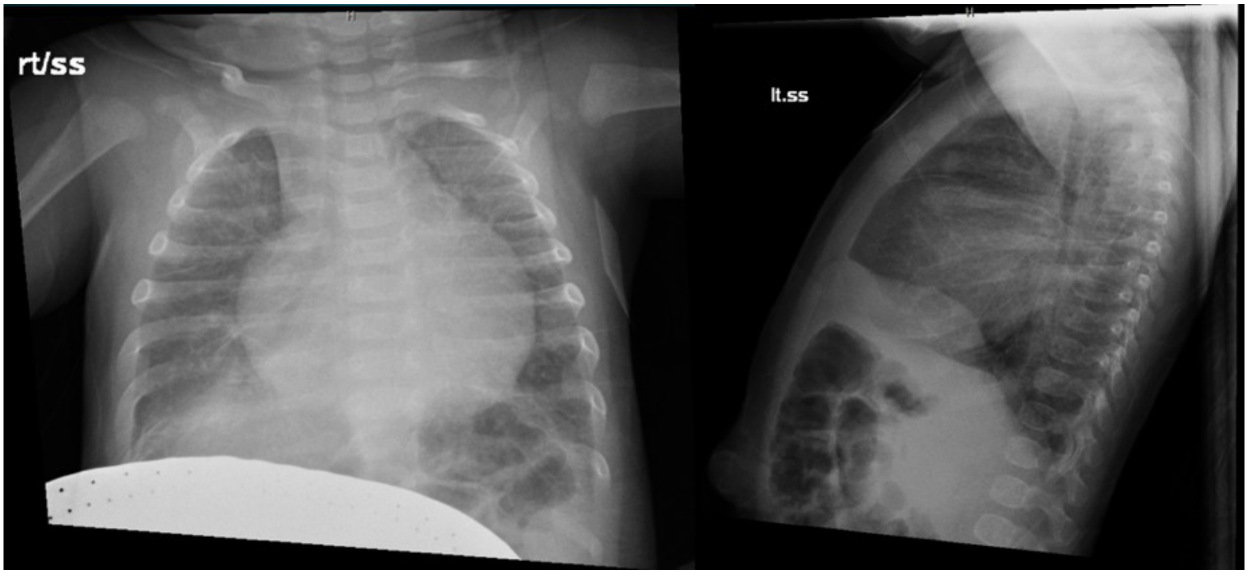


**Figure 1:** Chest radiograph in anteroposterior and lateral views showing cardiomegaly and pulmonary congestion on presentation at 3-months of age

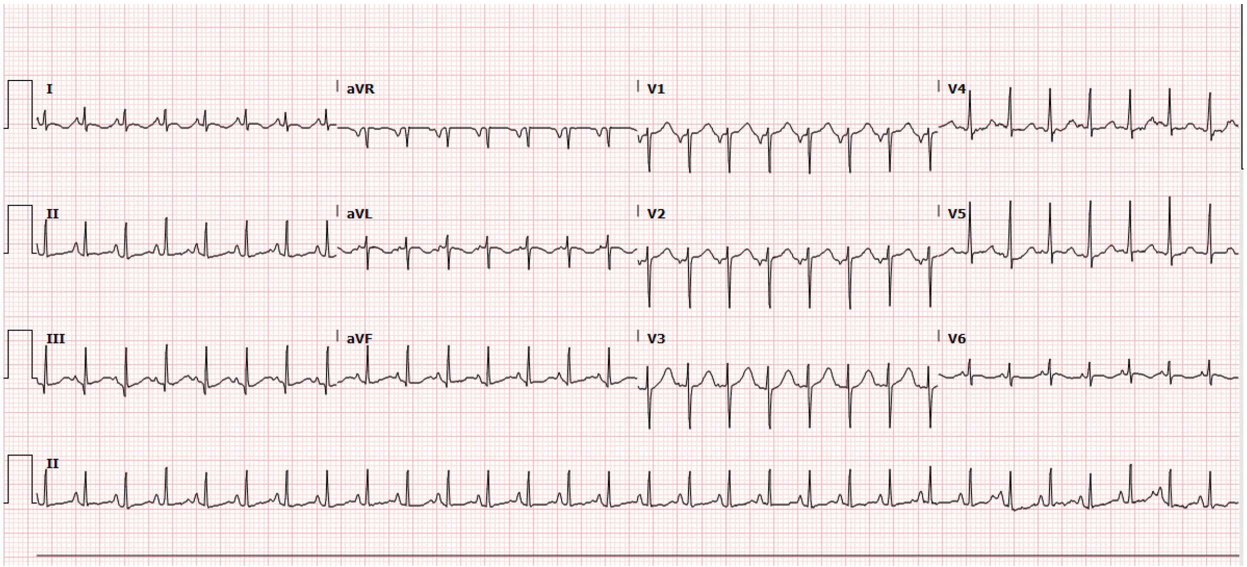


**Figure 2:** Electrocardiogram on presentation at 3-months of age

### Case 2: D308N/T340Nfs*25

Case 2 was transferred to our institution at day of life (DOL) 5 for heart failure evaluation. He was born to a 41-year-old G3P1011 mother at 38 weeks and 6 days gestation. His prenatal course was notable for fetal hydrops in the third trimester prompting a C-section. APGAR scores were 5 at one minute and 8 at 5 minutes of life. Birth weight was 4042 grams. He was intubated immediately for hypoxic respiratory failure. On DOL 4, a trial of extubation was unsuccessful, and a chest X-ray was notable for cardiomegaly. An echocardiogram showed severely depressed biventricular dysfunction (left ventricular ejection fraction 23%, normal 50%–70%) with a structurally normal heart. He was initiated on milrinone 0.5 mcg/kg/min and epinephrine at 0.4 mcg/kg/min and transferred to our institution for further management. We were able to extubate him on DOL 7 and wean him off epinephrine on DOL 9. However, he became tachycardic on DOL 14 with signs of decreased end-organ perfusion that required additional support with dobutamine at 3 mcg/kg/min. Echocardiogram at that time demonstrated a moderately dilated left atrium and severely dilated left ventricle, severely decreased left ventricular systolic function (ejection fraction 15%), moderate mitral valve regurgitation, and moderately reduced right ventricular systolic function. Rapid genome sequencing identified two *RPL3L* variants in trans (c.922G>A (p.Asp308Asn, D308N) & c.1018dup(Thr340Asnfs*25, T340Nfs*25). The proband's father is heterozygous for the D308N variant and the proband's mother is heterozygous for the T340Nfs*25 variant. Both parents are healthy, and both are of Hispanic and Caucasian descent with no consanguinity. Family history is unremarkable with a 6-year-old sister who is healthy. She had a normal cardiac evaluation but has not had genetic testing.

On DOL 22 the patient was put on the waitlist for a heart transplant. At the time he was on room air, tolerating enteral feeds with a heart failure regimen including dobutamine and milrinone infusions, digoxin, furosemide, and chlorothiazide. Over the next several weeks his clinical status deteriorated requiring escalation in respiratory support to positive pressure ventilation, with rising brain natriuretic peptide levels >70000 pg/nl (normal 37–1000 pg/nl) and increased inotropic and diuretic requirements. At five weeks of age, he underwent Berlin left ventricular assist device (LVAD) implant. His post-LVAD course was complicated by bilateral cerebral embolic infarcts and seizures, recurrent line infections, and a driveline site infection. Ultimately, he underwent orthotopic heart transplant with LVAD explant at seven months of age. His intraoperative course was notable for acute graft dysfunction requiring central venoarterial extracorporeal membrane oxygenation cannulation. He had significant improvement in cardiac function post-operatively and was decannulated on post-operative day (POD) 2, his chest was closed on POD 4, and he was successfully extubated on POD 5. He was discharged home on POD 19. Currently, he is doing well in the outpatient setting on an oral medical regimen of enalapril, amlodipine, and sildenafil, with an immunosuppression regimen of tacrolimus, mycophenolate, and prednisone. He is gaining weight and working with a nutritionist. A recent cardiac catheterization showed good hemodynamics, and an endomyocardial biopsy was negative for any evidence of rejection. He has no obvious neurologic sequelae, has been seizure-free, and is currently off all anti-epileptic medications.

### Case 3: D308N/c.1167+1G>A

Case 3 was transferred to our institution on day of life 2 for heart failure evaluation. She was the product of a naturally conceived pregnancy to a 27-year-old G1P0-P1 mother complicated by maternal rheumatoid arthritis treated with chloroquine and abnormal non-stress tests in the last 2 weeks of pregnancy. She was born at 38 weeks 6 days gestation weighing 3540g. APGARS were 5 at 1 minute and 8 at 5 minutes. She had poor respiratory effort at delivery requiring CPAP. She had worsening desaturation and increasing oxygen requirement on day of life (DOL) 1. Chest x-ray showed cardiomegaly and echocardiogram noted severe left ventricular and moderate right ventricular dysfunction, PDA with left-to-right shunting, and PFO with left-to-right shunting. She subsequently developed narrow pulse pressures, bradycardia, and hypoxemia and progressed to cardiac arrest. ROSC was achieved after 2 doses of epinephrine; she was intubated and transferred to our institution for further management. Left ventricular ejection fraction at time of admission to our institution was 42%. She was initiated on dopamine, epinephrine, and milrinone. Initial B Natriuretic Peptide was 13,958.3 pg/mL (normal 0-100 pg/mL) and troponin was 0.13 ng/mL. Metabolic and genetic work-up was initiated, including rapid exome sequencing with mitochondrial DNA analysis, which identified biparentally inherited variants in RPL3L. Both parents are healthy with no known cardiac disease, however family history was notable for a paternal first cousin once removed who died in infancy due to hypertrophic cardiomyopathy with skeletal muscle involvement.

Cardiac MRI on DOL17 showed severe left ventricular dilation and dysfunction, with LVEF of 25%, mild RV dilation, diffuse fibrosis and edema, and a type B aortic dissection. She had progressive left ventricular dysfunction and attempts at milrinone weans and extubation were unsuccessful.

She underwent Berlin EXCOR left ventricular assist device (LVAD) on DOL24 and was listed for heart transplant at 7 weeks of age. Her clinical course while awaiting transplant was complicated by medical necrotizing enterocolitis, feeding intolerance requiring TPN and feed optimization, upper GI bleeding secondary to multiple ulcerations, cholelithiasis requiring ursodiol, occlusive thrombus in right CFA requiring lovenox, seizures in the setting of multiple right MCA strokes while on LVAD. She underwent heart transplant at 9 months of age which without major complications. She was 21 months of age at most recent follow-up and currently doing well in the outpatient setting on oral immunosuppression with no evidence of rejection to date. She is off antiepileptic medication, seizure-free, and developing appropriately for age.

**Supplementary Table 1: Oligos Used**

| Oligo | Sequence Forward | Sequence Reverse |
|---|---|---|
| RPL3 shRNA | AGACGCTAGCCCTATCTGACAAGAG CATCAAAACTAGTATTGATGCTCTTG TCAGATAGGTTTTTGAATTCT | AGAATTCAAAAACCTATCTGACAAG AGCATCAATACTAGTTTTGATGCTC TTGTCAGATAGGGCTAGCGTCT |
| RPL3 qPCR | TGAAGAGCTTCCCTAAGGATGA | CTTCCCGCACGATGTGAGTC |
| RPL3L qPCR | CCCCACTACGGGGAAGTGA | GAGGGACTTTCTCAGCGTAATG |
| GAPDH | TGATGACATCAAGAAGGTGGTGAAG | TCCTTGGAGGCCATGTGGGCCAT |
| RPL3 Promoter 1 | ACGTCAACTTCTGAACGAAAG | CGGGTCCGCTATATAAAGCCA |
| RPL3 Promoter 2 | GGAAGAGCGTGCGTGGAAATG | CCATCAAATCCCGCCGGTAGA |
| RPL3 Upstream 1 | ACAGAGCGAGACTCCGTCTCA | AACGGCCTCACGACCTCTGGT |
| RPL3 Upstream 2 | ATCTCAGGTGAACCACCCAC | GAGATCGTGCCATTGTGCTC |
| RPL3 Upstream 3 | TCTTGGCTTACTGCAACCTCC | ACCTGAGATCAGGAGTTCGAGA |
| RPL4 Promoter 1 | TCCGTGTTGGTTCTTAGGCG | TGCTGCCACAGGAAAAGGAA |
| RPL4 Promoter 2 | GGAAGGTGACATATACAGCGGG | GAGAGAGGAGACAGCCACGC |
